# Effect of a Growth Factor-Rich Platelet Lysate Gel with or without *Uncaria tomentosa* Extract on a Rat Model of Postmenopausal Vaginal Atrophy

**DOI:** 10.1101/2025.09.19.677269

**Authors:** Diego Hidalgo-Avendaño, Christian Roberto Pitot Alvarez, Shaloom Samantha Hurtado Ollero, Ximena Flores Velasquez, Angie Surco Quispe, Arianna Marín Gonzales, José Aguilar-Olano

**Affiliations:** School of Medicine, Universidad Peruana Cayetano Heredia, Lima, Peru; School of Veterinary Medicine and Animal Science, Universidad Peruana Cayetano Heredia, Lima, Peru; Graduate School, Universidad Peruana Cayetano Heredia, Lima, Peru; School of Science and Engineering, Universidad Peruana Cayetano Heredia, Lima, Peru

**Keywords:** Vaginal atrophy, Platelet lysate, Uncaria tomentosa, Tissue repair, anti inflammatory

## Abstract

The climacteric and menopausal transitions involve significant hormonal and physiological changes that impact women’s health. During the postmenopausal period, various changes have been documented, including atrophy of the vaginal mucosa and a chronic low-grade inflammatory state, particularly affecting the vaginal mucosa, which contributes to symptomatic deterioration and reduced quality of life. In this study, a chitosan-based polymeric gel was used as a delivery matrix for natural compounds, such as growth factors derived from platelet-rich plasma (PRP) and a hydroalcoholic extract of *Uncaria tomentosa* (UG), for intravaginal application in a menopausal animal model with vaginal atrophy. A 20% concentration of platelet lysate (PL), with or without UG, showed comparable outcomes to estrogen treatment, including restoration of the squamous epithelium, increasing the number of epithelial layers in the vaginal lining with histological evidence of absent chronic inflammatory infiltration. In contrast, 30% PL alone promoted epithelial repair without inflammation; however, when combined with UG, it failed to support tissue repair and was associated with chronic inflammation. Our findings suggest that a 20% platelet lysate formulation may effectively reverse vaginal atrophy without inducing adverse inflammatory responses. This formulation may represent a viable non-hormonal therapeutic option for managing postmenopausal vaginal atrophy to improve the clinical and histological characteristics of postmenopausal vaginal tissues.

## Introduction

Menopause marks the end of a woman’s reproductive stage and is clinically defined as the permanent cessation of menstruation due to a decline in estrogen levels. The subsequent stage, postmenopause, is a vital period that comprises nearly 40% of a woman’s life. This period is associated with a heightened risk of clinical symptoms (1). Among these symptoms are hot flashes, mood disturbances, and most notably, vaginal atrophy, which significantly impairs quality of life (2).

To address postmenopausal symptoms, hormone replacement therapy (HRT) has been developed using conventional hormones or synthetic estrogen-like compounds that help alleviate symptoms of vaginal atrophy. While these drugs have succeeded in reducing some complications, they have not been effective for all postmenopausal conditions. Moreover, in some cases, their use is limited by adverse effects and high cost, resulting in persistent discomfort and diminished quality of life (3).

During menopause, hormonal changes such as decreased estrogen levels and increased follicle-stimulating hormone (FSH) levels are well recognized. In addition, elevated levels of pro-inflammatory cytokines—such as IL-1, IL-6, and tumor necrosis factor-alpha (TNF-α)—have been reported, contributing to a state of low-grade chronic inflammation (4). Other systemic changes also occur, including vascular tone loss and increased expression of adhesion molecules in endothelial cells (5). Although these changes affect the body systemically, they also manifest locally. For instance, chronic inflammation has been observed in the vaginal epithelium, contributing to the aforementioned symptoms (6). Thus, reducing chronic inflammation could improve vaginal epithelial integrity and overall clinical outcomes.

Current research focuses on identifying non-hormonal alternatives with anti-inflammatory and regenerative properties to address symptoms refractory to HRT. This study proposes the use of compounds with potential anti-inflammatory effects—such as gel-forming polymers, platelet-rich plasma (PRP), and *Uncaria tomentosa* (UG)—to reduce vaginal atrophy.

The selected polymer must be compatible with the unique conditions of the vaginal cavity, resistant to pH fluctuations, exhibit high mucoadhesiveness, facilitate mucosal penetration for adequate growth factor delivery, and ensure stability in vaginal fluids. Among the widely studied polymers, chitosan—a derivative of the natural polymer chitin—was chosen for this study due to its favorable properties (7).

PRP has recently gained interest in gynecology, showing promising results in applications such as vulvar reconstruction, wound healing, and treatment of skin lesions (8). In rat models, autologous PRP has demonstrated effectiveness in cutaneous wound healing and endometrial regeneration (9). Favorable outcomes have also been reported with human-derived PRP applied to rat models (10). Given its documented efficacy in tissue regeneration and wound healing, PRP was incorporated as the primary bioactive component of the formulation for the gel, and platelet lysate (PL)—containing released growth factors—was used to avoid cross-species incompatibility.

*Uncaria tomentosa* (UG) has been used for various conditions due to its antimutagenic, antiviral, antioxidant, and anti-inflammatory properties (11). It contains several secondary metabolites, including pentacyclic alkaloids, which exhibit anti-inflammatory, pro-apoptotic, and immunomodulatory effects (12). Its immunomodulatory activity is mediated by inhibition of pro-inflammatory cytokines such as TNF-α and IL-6, along with modulation of IL-10 (13,14). Additionally, UG has been shown to inhibit the activation of transcription factors such as NF-κB (15), which helps to reduce tissue inflammation. Given its capacity to suppress key inflammatory mediators, *Uncaria tomentosa* was included in the gel formulation.

This study aims to evaluate the effect of combining growth factors derived from PRP lysate, with or without a standardized extract of *Uncaria tomentosa*, on clinical outcomes and vaginal histology in a postmenopausal rat model.

## Materials and Methods

### Methodological Design

Our experimental study was conducted using a postmenopausal vaginal atrophy animal model with Sprague Dawley rats (16). The sample size was calculated based on previously reported efficacy rates: 86.7% for platelet-rich plasma (PRP) and 51.7% for *Uncaria tomentosa,* assuming a combined expected efficacy of 95%. A power analysis was performed using these estimates, with a confidence level of 95% and a statistical power of 80%, resulting in a minimum required sample size of nine animals per group (Table 1).

**Table 1:** Distribution of experimental groups.

| GROUP | SAMPLE | TREATMENT |
| --- | --- | --- |
| WHITE CONTROL (CO) | 9 | Healthy rats without experimental testing. |
| NEGATIVE CONTROL (C1) | 9 | Untreated ovariectomized rats |
| POSITIVE CONTROL (C2) | 9 | Ovariectomized rats with topical estrogen treatment |
| TREATED GROUP A | 9 | Ovariectomized rats treated with 20% PL |
| TREATED GROUP B | 9 | Ovariectomized rats treated with 20% PL and 200 ug/ml UG |
| **TREATED GROUP C** | 9 | Ovariectomized rats treated with 30% PL |
| **TREATED GROUP D** | 9 | Ovariectomized rats treated with 30% PL and 200 ug/ml UG |

Note: PL = Platelet lysate; UG = *Uncaria tomentosa*.

### Animals

Female Sprague Dawley rats aged 3 months (∼260 g) were used, housed under controlled conditions, and fed with phytoestrogen-free rat chow. They were randomly assigned to experimental groups. At the end of the study, euthanasia was performed using sodium pentobarbital (17). The study was approved by the Animal Ethics Committee of the Universidad Peruana Cayetano Heredia (UPCH, code 210380).

### Platelet-Rich Plasma

PRP was obtained from six healthy voluntary donors at the Blood Bank of the National Hospital Cayetano Heredia (HNCH), who signed informed consent. The PRP was then used to prepare PL through freeze-thaw cycles, centrifugation, and the addition of 10% calcium gluconate. The PL was filtered (0.22 µm) and aliquoted according to groups.

### Hydroalcoholic Extract

Dry powder derived from the bark of Uncaria tomentosa (code 6.999) was generously provided by Dr. Mirtha Navarro (Costa Rica). The drying technique, previously described using methanol (MeOH), was followed as outlined in reference (18). The powder was reconstituted in 20° ethanol, and polyphenolic compounds were quantified by High-Performance Liquid Chromatography (HPLC).

### Chitosan

Chitosan powder, donated by Dr. José Bauer (UPCH), was extracted from squid pens and characterized by a deacetylation degree of 86% and a moisture content of 4.223%. The powder was ground using an analytical mill and subsequently sieved through a W.S. Tyler stainless steel mesh No. 60.

### Gel

A 2.4% chitosan gel was formulated in 2% acetic acid, with phenoxyethanol added as a preservative agent (19). The mixture was sterilized in an autoclave (20), then the *Uncaria tomentosa* extract and PL (20% or 30%) were added under sterile conditions inside a laminar flow cabinet. The gel was incubated at 37°C until treatment completion.

### Surgical Menopause Animal Model

Bilateral ovariectomy was performed on all rats except the positive control group, following the Abramov protocol (21). Anesthesia was induced with ketamine and xylazine, and pre– and postoperative care was provided, including analgesia (tramadol) and antibiotics (enrofloxacin) for four days.

### Direct Vaginal Cytology

Vaginal lavage samples were collected before surgery to confirm the estrous cycle and four weeks post-surgery to verify the absence of estrus phase. Samples were microscopically analyzed to determine the estrous cycle phases (22).

### Treatment Administration

Ten days after ovariectomy, the prepared gels were applied intravaginally (0.4 mL) once daily for 10 consecutive days using a modified pipette.

### In Vitro Measurement of PDGF-BB Concentration

Hydrogels from each experimental group were incubated in phosphate-buffered saline (PBS, pH 7.4), and PDGF-BB release was assessed at 24, 48, 96, 144, 192, and 240 hours (Table 2). PDGF-BB levels were quantified using the Human PDGF-BB ELISA Kit (ELK2289). Supernatants were collected at each time point, and ELISA plates were prepared with standards, blanks, and samples. The assay followed the standard ELISA protocol, including sequential incubations, washing steps, addition of biotinylated antibodies, streptavidin-HRP, TMB substrate, and stop solution. Absorbance was measured at 450 nm using a microplate reader.

**Table 2:** Distribution of groups in vitro.

|  |  |
| --- | --- |
| **Group 1** | Gel with 20% PL |
| **Group 2** | Gel with 30% PL |
| **Group 3** | Gel with 20% PL and 200ug/ml UG |
| **Group 4** | Gel with 30% PL and 200ug/ml UG |
| **Group 5** | PL at 20% |
| **Group 6** | PL at 30% |
| **Group 7** | Gel |
| **Group 8** | Gel + UG 200 ug/ ml |

Note: *Q* = Chitosan; *PL* = Platelet lysate; *UG* = *Uncaria tomentosa*.

### Histological Study of Vaginal Tissue Post-Treatment

Following euthanasia with pentobarbital sodium (17), vaginal tissues were collected, fixed in 4% paraformaldehyde, dehydrated through a graded ethanol series, embedded in paraffin, and sectioned at 4–6 µm thickness. Tissue sections were stained with hematoxylin and eosin (H&E) and Masson’s trichrome, and examined under light microscopy at 10× to 40× magnification. Histological parameters—including epithelial thickness, presence of leukorrhea, inflammatory infiltrates, and re-epithelialization—were assessed and compared across the control groups (C0, C1, and C2) (Fig 1).

**Fig 1.**
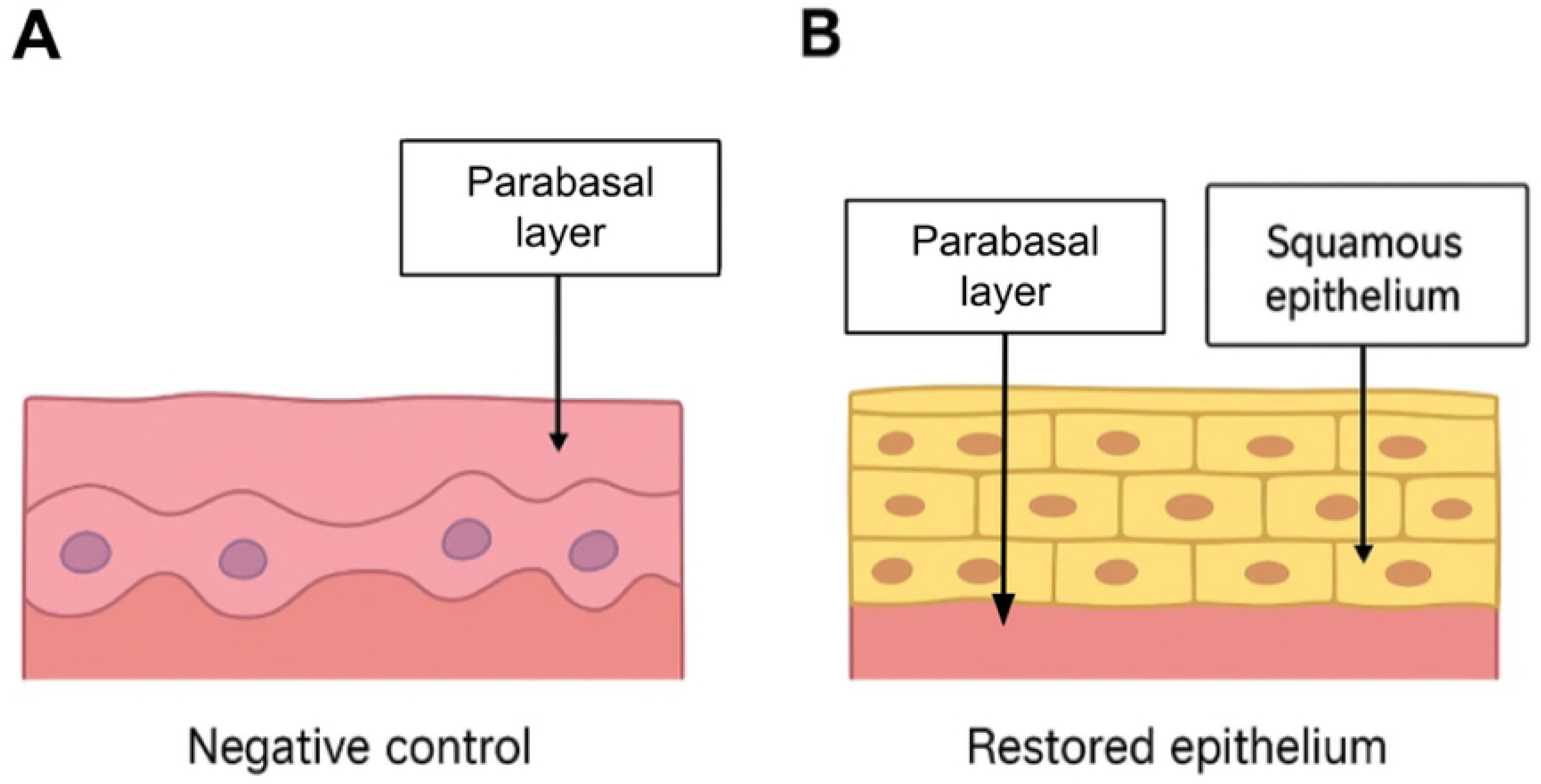
Comparison of the vaginal lining epithelium. A = Negative control, B = Restored epithelium. In A, the squamous epithelium is absent and only the parabasal layer is present; in B, squamous epithelium is visible.

### ELISA Study of Estradiol Concentration

Estradiol levels were measured before ovariectomy and prior to euthanasia by collecting blood from the tail vein. The Rat E2 ELISA kit (ELK8714) was used. The standard ELISA procedure was followed: preparation of wells for standards and blanks, addition of samples and biotinylated conjugate, incubations, washes, streptavidin-HRP application, TMB substrate, and stop reagent. Absorbance was measured at 450 nm.

### Ethical Considerations

The study was approved by three ethics committees: the Institutional Ethics Committee for Human Research and the Animal Use Ethics Committee of Universidad Peruana Cayetano Heredia (protocol number 210380), as well as the Research Ethics Committee of the National Hospital Cayetano Heredia (approval code No. 48, 2023).

### Statistical Analysis

Statistical tests were chosen according to the variable type: Kruskal-Wallis for non-parametric qualitative data, one-way ANOVA for parametric data, and chi-square test for contingency tables. A p-value < 0.05 was considered statistically significant. Analyses were performed using SPSS versions 25 and 29.

## Results

### Vaginal Lavage

#### Preoperative Period

Vaginal lavages were performed for five consecutive days, two weeks prior to surgery. The presence of keratinized, nucleated epithelial cells and leukocytes allowed determination of the estrous cycle phase. All rats were confirmed to be in estrus, indicating active ovarian function.

#### Postoperative Period

Four weeks after ovariectomy, a marked reduction in nucleated and squamous epithelial cells was observed in all groups, except for the operated control group (C0), suggesting decreased estrogen levels due to ovariectomy.

### PDGF-BB Quantification

#### Calibration Curve

A calibration curve was obtained with a determination coefficient (R²) close to 1, confirming measurement accuracy.

#### PDGF-BB Release Profiles

Groups containing 20% or 30% PL demonstrated sustained PDGF-BB release, maintaining concentrations above 3000 pg/mL for up to 144 hours before gradually declining. In contrast, groups containing chitosan alone or chitosan combined with Uncaria tomentosa exhibited negligible PDGF-BB release (Table 3, Fig 2).

**Fig 2.**
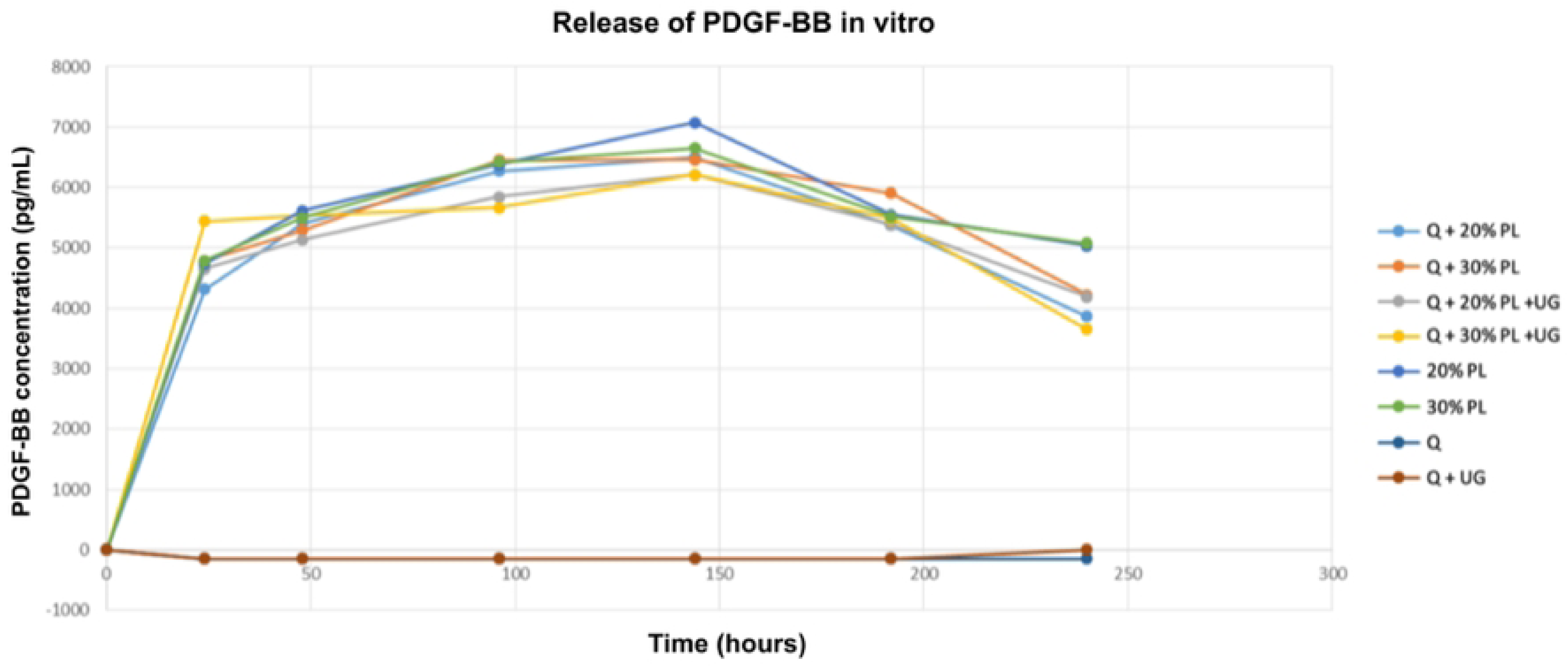
Release of PDGF-BB in vitro. Hydrogels from each experimental group were incubated in PBS, and PDGF-BB release was assessed at 24, 48, 96, 144, 192, and 240 hours by ELISA technique. Q = Chitosan; PL = Platelet lysate; UG = Uncaria tomentosa.

**Table 3:** PDGF-BB concentrations of the different groups.

| Time<br>(hrs) | Q + 20%<br>PL | Q + 30%<br>PL | Q + 20% PL<br>+ UG | Q + 30% PL<br>+ UG | 20% PL | 20% PL | Q | Q + UG |
| --- | --- | --- | --- | --- | --- | --- | --- | --- |
| 24 | 4310,417 | 4785,625 | 4653,792 | 5444,583 | 4737,250 | 4783,875 | -145,458 | -146,708 |
| 48 | 5411,625 | 5295,417 | 5132,458 | 5532,583 | 5619,125 | 5499,583 | -145,583 | -146,375 |
| 96 | 6270,375 | 6450,708 | 5849,750 | 5661,542 | 6374,417 | 6415,750 | -149,458 | -147,208 |
| 144 | 6502,833 | 6465,133 | 6211,708 | 6209,750 | 7077,458 | 6651,625 | -147,333 | -147,917 |
| 192 | 5378,917 | 5909,583 | 5381,875 | 5479,792 | 5551,542 | 5519,667 | -147,625 | -146,417 |
| 240 | 3867,000 | 4224,000 | 4189,458 | 3650,542 | 5036,333 | 5079,125 | -145,792 | -146,250 |

Note: Q = Chitosan; PL = Platelet lysate; UG = *Uncaria tomentosa*.

### Phytochemical Composition of the Hydroalcoholic Extract of *Uncaria tomentosa*

Five polyphenols were identified in the extract, including procyanidin and propelargonidin dimers, as well as procyanidin trimers. The most abundant compound was a procyanidin dimer, accounting for 41.3% of total content, with a concentration of 89.15 mg/g higher than previously reported values (23). Total phenolic content measured by the Folin–Ciocalteu method was 359 mg GAE/g of dry extract, consistent with earlier studies (23).

The ORAC assay revealed an antioxidant capacity of 5.6 mmol TE/g, in line with prior reports (18). The IC50 was 6.21 mg extract/L, indicating stronger antioxidant activity compared to similar extracts reported in the literature (e.g., 13.5 mg/L) (24).

### External Morphology of Vulva and Uterus

Gross examination of the vulvar and uterine tissues revealed no statistically significant differences in external morphology between treatment and control groups. All exhibited a reddish, dry vulva with a contracted opening, except for group C0, which showed a dilated vaginal opening during estrus.

### Estradiol Measurement

#### Calibration Curve

The standard curve showed a second-degree polynomial regression with an R² = 0.99, indicating high linearity and reliability of the assay.

#### Preoperative and Postoperative Estradiol Concentration

Before surgery, peripheral blood estradiol levels ranged from 22 to 49 pg/mL across all groups, consistent with normal estrous cycles. Following ovariectomy, estradiol levels decreased in all groups except the control (C0) and the group treated with topical estradiol (C2) (Fig 3).

**Fig 3.**
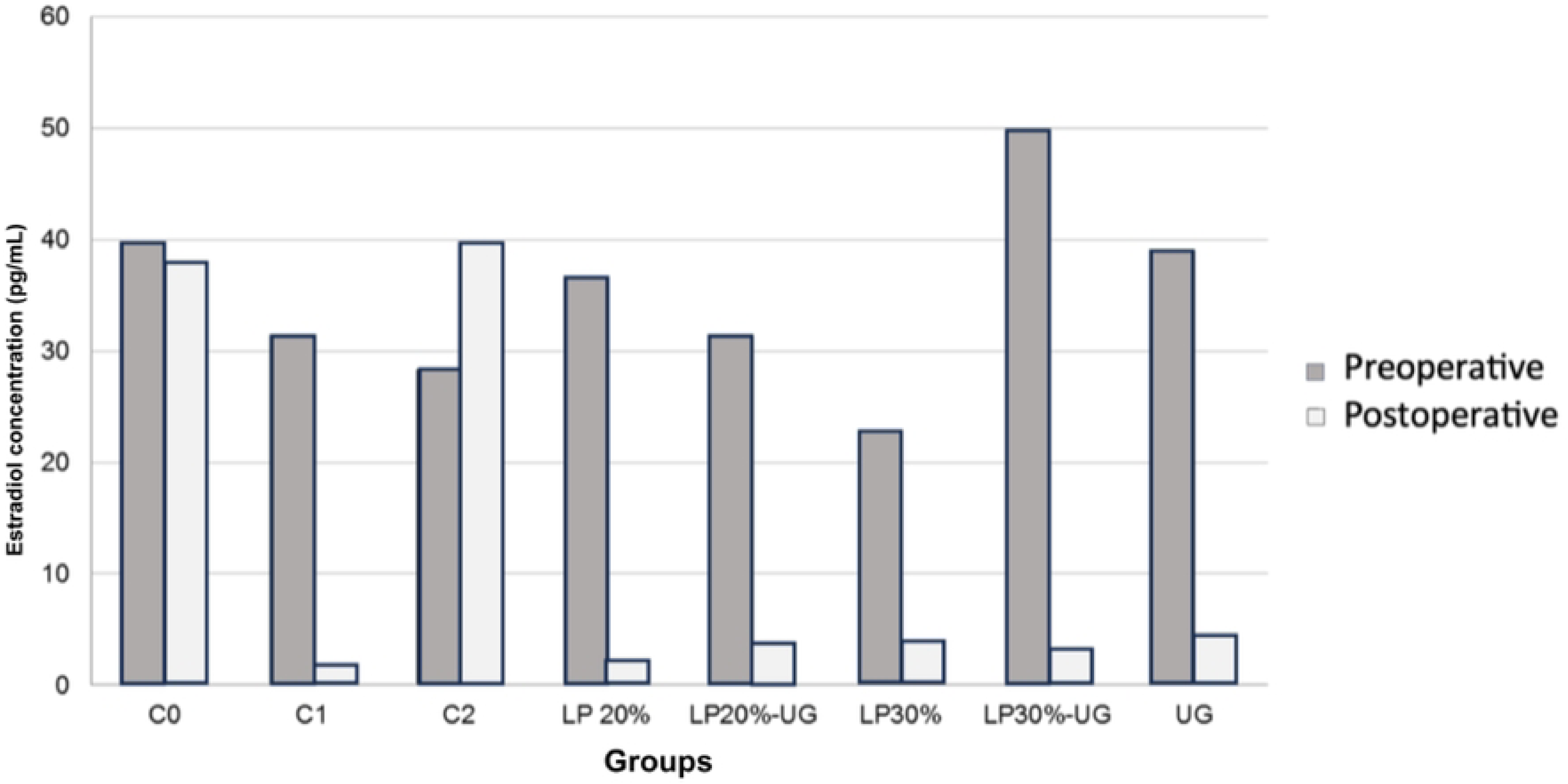
Estradiol concentrations before and after oophorectomy. C0 = Control without operation, C1 = Control without treatment, C2 = Control with topical estrogen. *Values represent measured estradiol concentrations (pg/mL); no statistical tests were performed*.

### Histological analysis

#### Vaginal Inflammation

The group treated with 30% PL plus cat’s claw extract (PL 30%-UG) exhibited significantly greater chronic inflammation compared to control groups C0 and C2 (p = 0.003 and p = 0.005, respectively) (Table 4). As shown in Fig 4, approximately 55% of the PL 30%-UG group presented chronic inflammation. In contrast, the other groups demonstrated no significant differences and approximately 65% of cases showed no signs of vaginal inflammation.

**Fig 4.**
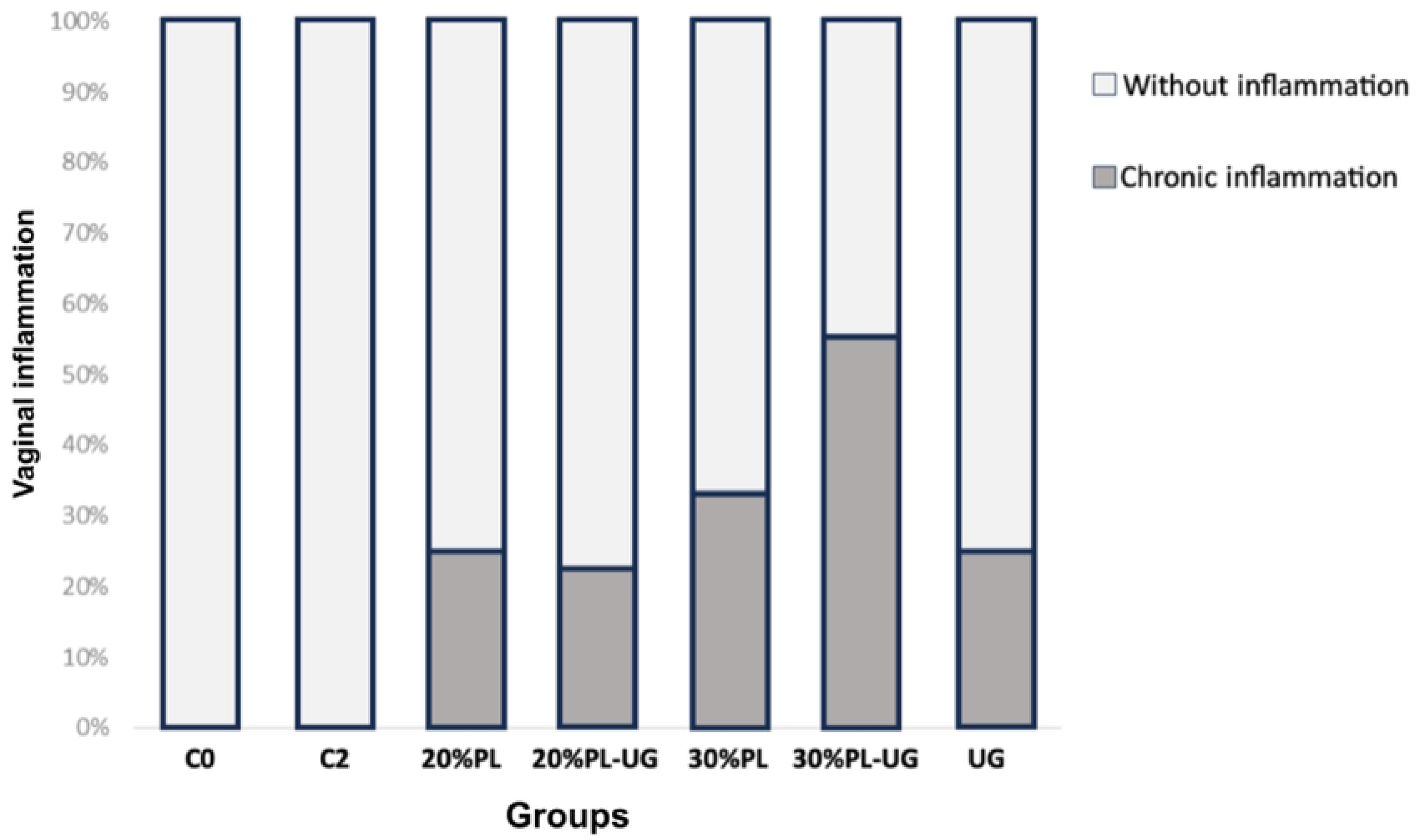
Percentages of cases with chronic inflammation or without vaginal inflammation. C0= Control without operation, C2= Control with estrogen. *Data represent observed proportions within each group; no statistical analysis was performed*.

**Table 4:**
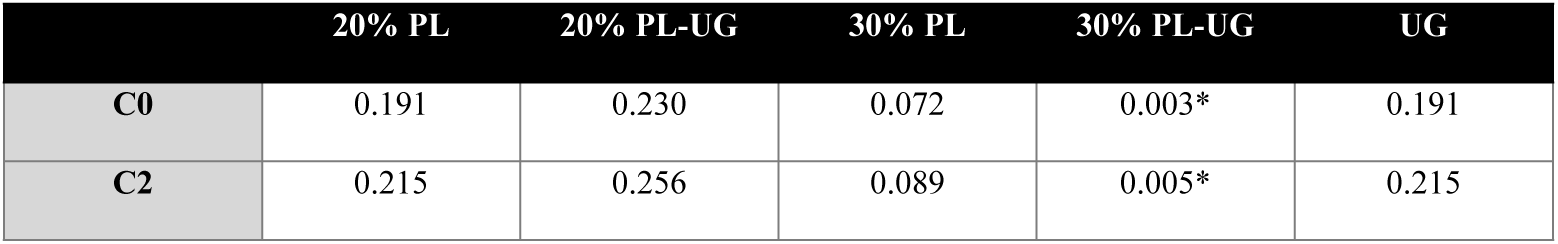
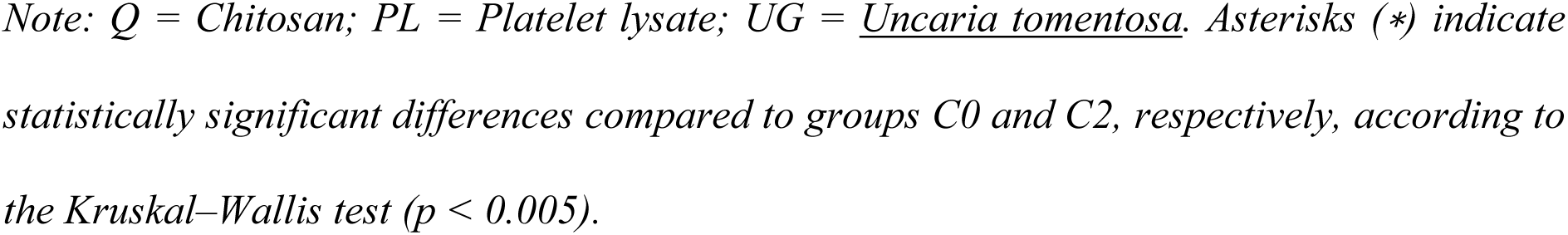
Comparison of vaginal inflammation between C0 and C2.

|  | 20% PL | 20% PL-UG | 30% PL | 30% PL-UG | UG |
| --- | --- | --- | --- | --- | --- |
| C0 | 0.191 | 0.230 | 0.072 | 0.003* | 0.191 |
| C2 | 0.215 | 0.256 | 0.089 | 0.005* | 0.215 |

Note: Q = Chitosan; PL = Platelet lysate; UG = *Uncaria tomentosa*. Asterisks (\*) indicate statistically significant differences compared to groups C0 and C2, respectively, according to the Kruskal–Wallis test ( $p < 0.005$ ).

#### Uterine Inflammation

No inflammation was detected in the uterus in any of the groups.

#### Vaginal Epithelium

Analysis revealed significant differences in vaginal epithelial types between the non-ovariectomized control group (C0) and the PL30%-UG and UG groups (p = 0.002 and p < 0.001, respectively) (Table 5). In the PL30%-UG and UG groups, absence of squamous epithelium predominated, with 66.6% and 87.5% of cases presenting parabasal flat epithelium, respectively (Fig 5). The C2 group, treated with estrogen, showed restoration of the squamous epithelium. Conversely, the C1 group (ovariectomized without treatment) exhibited parabasal flat epithelium but differed significantly from the PL20%, PL20%-UG, and PL30% groups, which preserved squamous epithelium (p = 0.020, p = 0.014, and p = 0.049, respectively) (Table 5).

**Fig 5.**
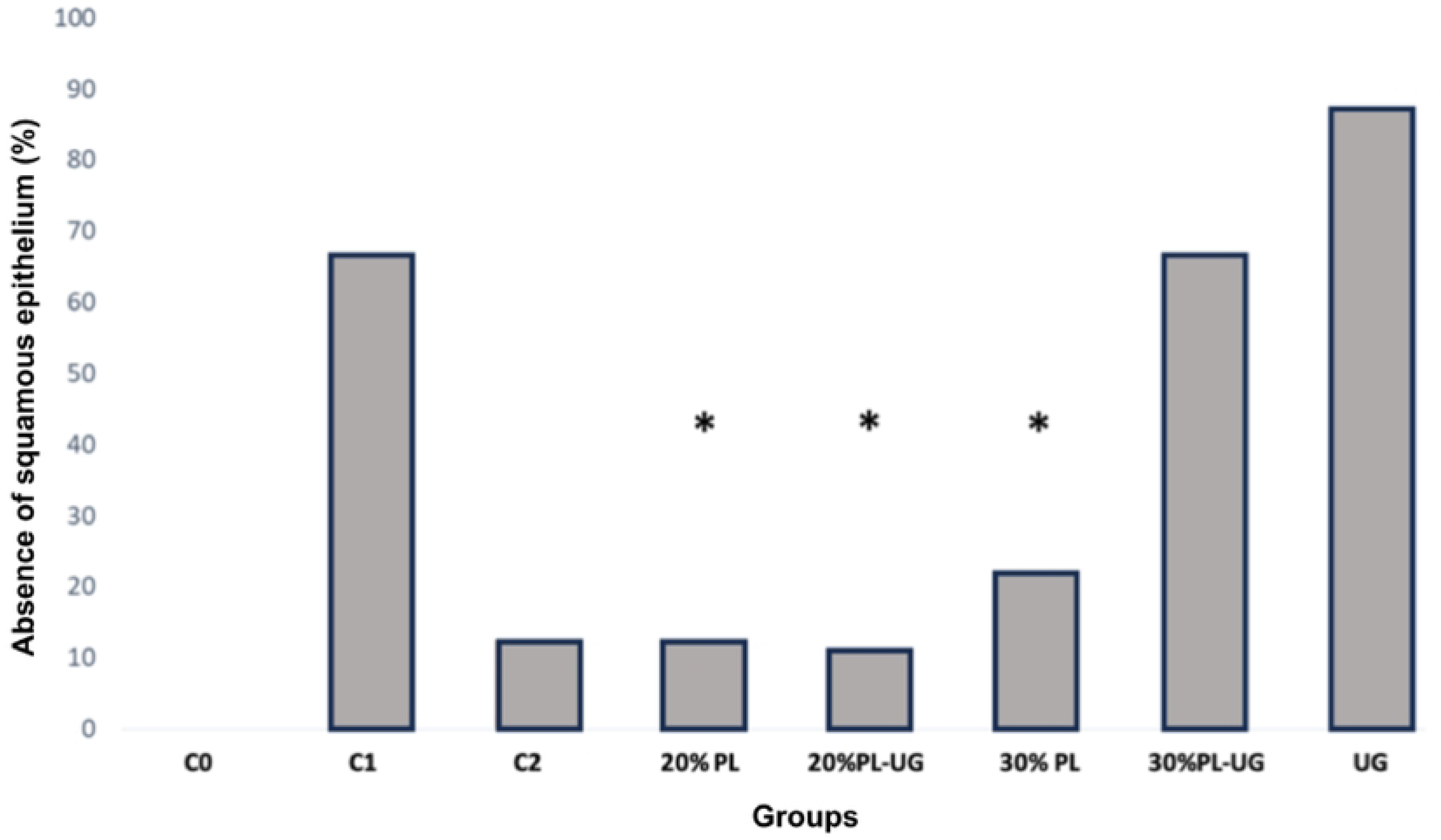
Percentage of cases with absence of squamous epithelium. C0 = Control without operation, C1 = Control without treatment, C2 = Control with estrogen. *Data are presented as percentages. Asterisks (∗) indicate statistically significant differences compared to C1 (p < 0.005)*.

**Table 5:**
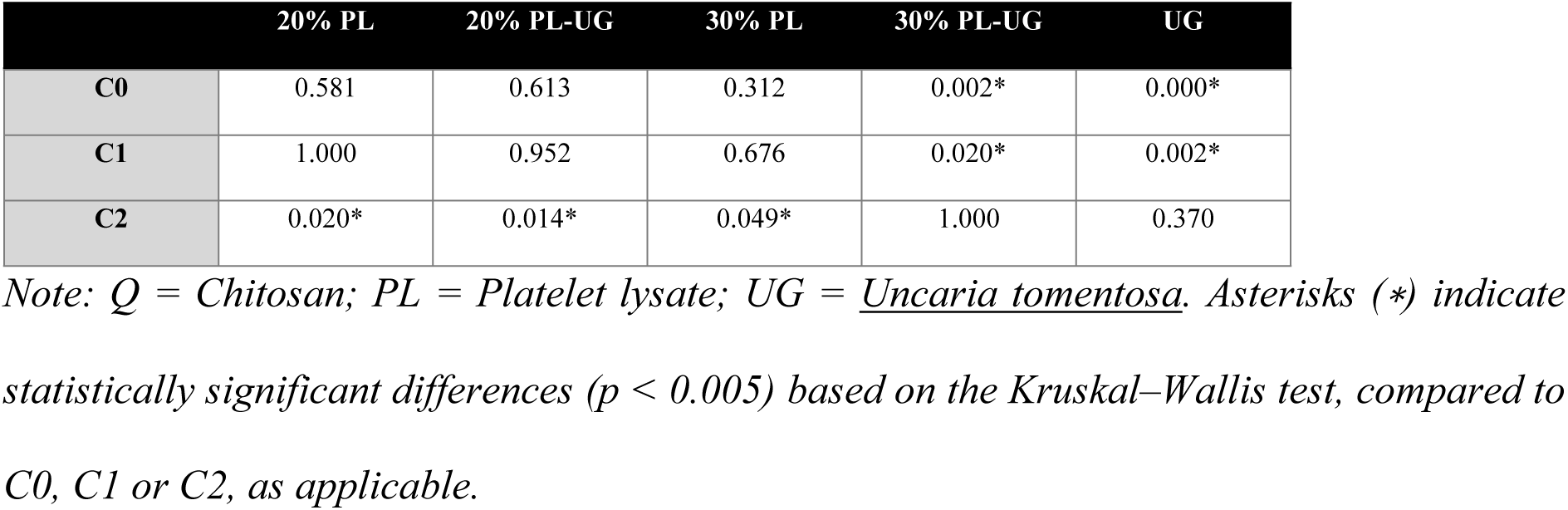
Comparison of vaginal lining epithelium with the control group.

#### Number of Vaginal Epithelium Layers

Groups C0 and C2 exhibited a median of 11 to 12 vaginal epithelial layers, with a range of 9 to 14 layers and a symmetrical distribution. The PL20% group showed a median of 8 to 9 layers with a wider range (3 to 14), indicating the highest variability among groups. Both PL20%-UG and PL30% demonstrated medians of 9 to 10 layers; PL20%-UG displayed a more concentrated distribution except for two outliers, whereas 75% of PL30% cases had between 5 and 10 layers, with the remaining 25% ranging from 10 to 12 layers. The PL30%-UG group had 75% of cases with 3 to 8 layers and 25% with 8 to 10 layers. Groups UG and C1 presented medians of 5 to 6 layers, with UG showing a more concentrated distribution compared to C1 (Fig 6).

**Fig 6.**
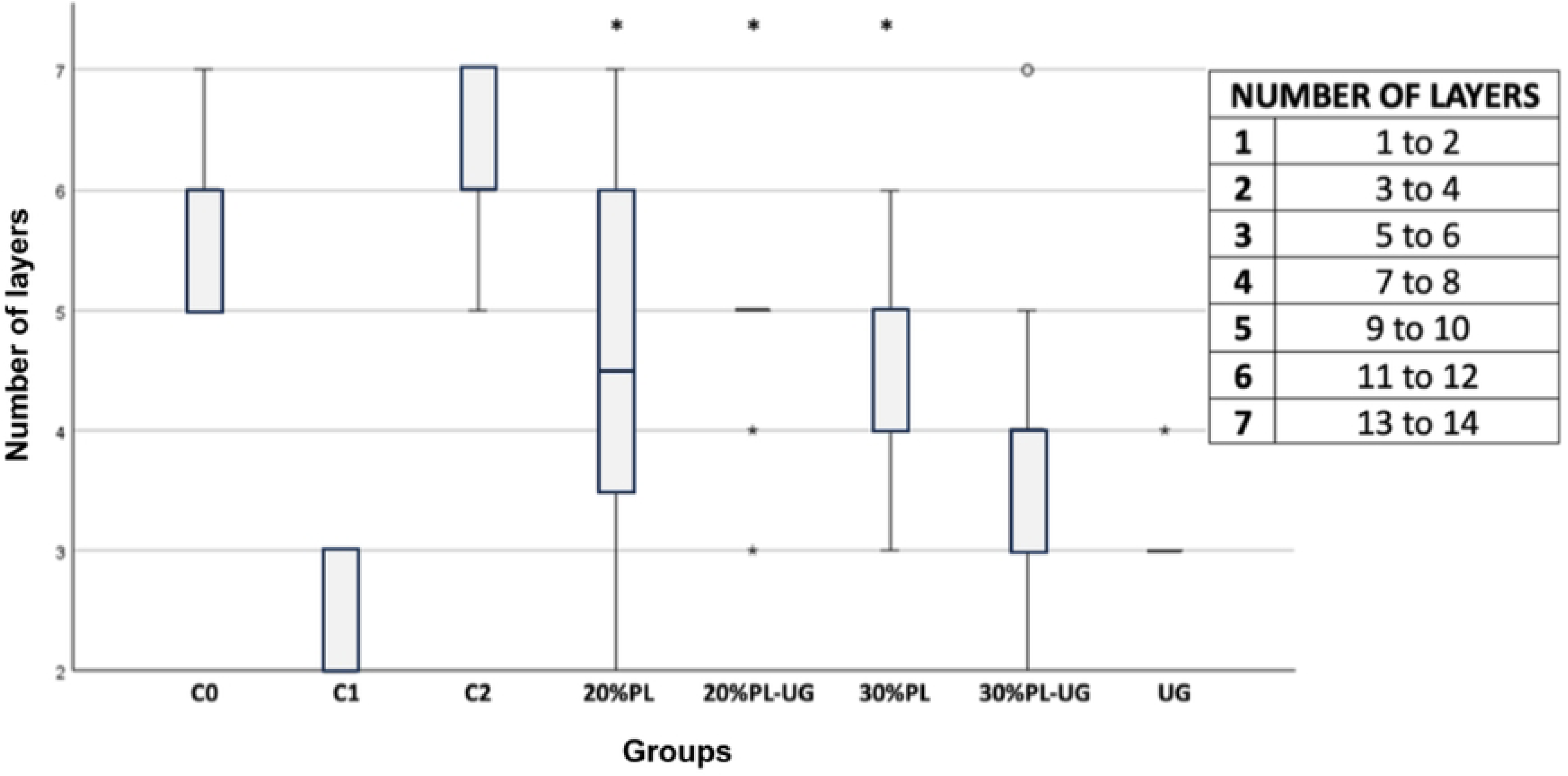
Number of layers of the vaginal lining epithelium. C0 = Control without operation, C1 = Control without treatment, C2 = Control with topical estrogen. *Values are presented as median and interquartile range. Asterisks (∗) indicate statistically significant differences compared to C1 (p < 0.005)*.

Statistical analysis revealed a significant decrease in the number of epithelial layers in the UG group compared to C0 and C2 (p = 0.003 and p = 0.001, respectively), while C2 differed significantly from PL30%-UG (p = 0.047). Additionally, the PL20%, PL20%-UG, and PL30% groups exhibited a significant increase in the number of layers compared to C1 (p = 0.016, p = 0.027, and p = 0.027, respectively) (Table 6).

**Table 6:** Comparison with controls by number of layers.

|  | 20% PL | 20% PL-UG | 30% PL | 30% PL-UG | UG |
| --- | --- | --- | --- | --- | --- |
| C0 | 1.000 | 1.000 | 1.000 | 0.189 | 0.003* |
| C1 | 1.000 | 0.886 | 0.886 | 0.047* | 0.001* |
| C2 | 0.016* | 0.027* | 0.027* | 0.708 | 1.000 |

Note: Q = Chitosan; PL = Platelet lysate; UG = *Uncaria tomentosa*. Asterisks (\*) indicate statistically significant differences based on the Kruskal–Wallis test ( $p < 0.005$ ) compared to C0, C1, or C2, as appropriate.

### Relationship Between Histological Variables

#### Association Between Vaginal Inflammation and Epithelial Lining

Table 7 shows that 71.4% of rats exhibited squamous epithelium and concurrently showed no signs of vaginal inflammation. A corrected residual greater than 1.96 indicates a statistically significant association. Fig 7 further illustrates the marked difference in the number of rats without inflammation that presented squamous epithelium. These findings confirm a significant relationship between the presence of squamous epithelium and the absence of vaginal inflammation (p = 0.013) (Table 8). Moreover, odds ratio analysis indicated that rats without inflammation were 4.5 times more likely to exhibit squamous epithelium in the vaginal lining (Table 9).

**Fig 7.**
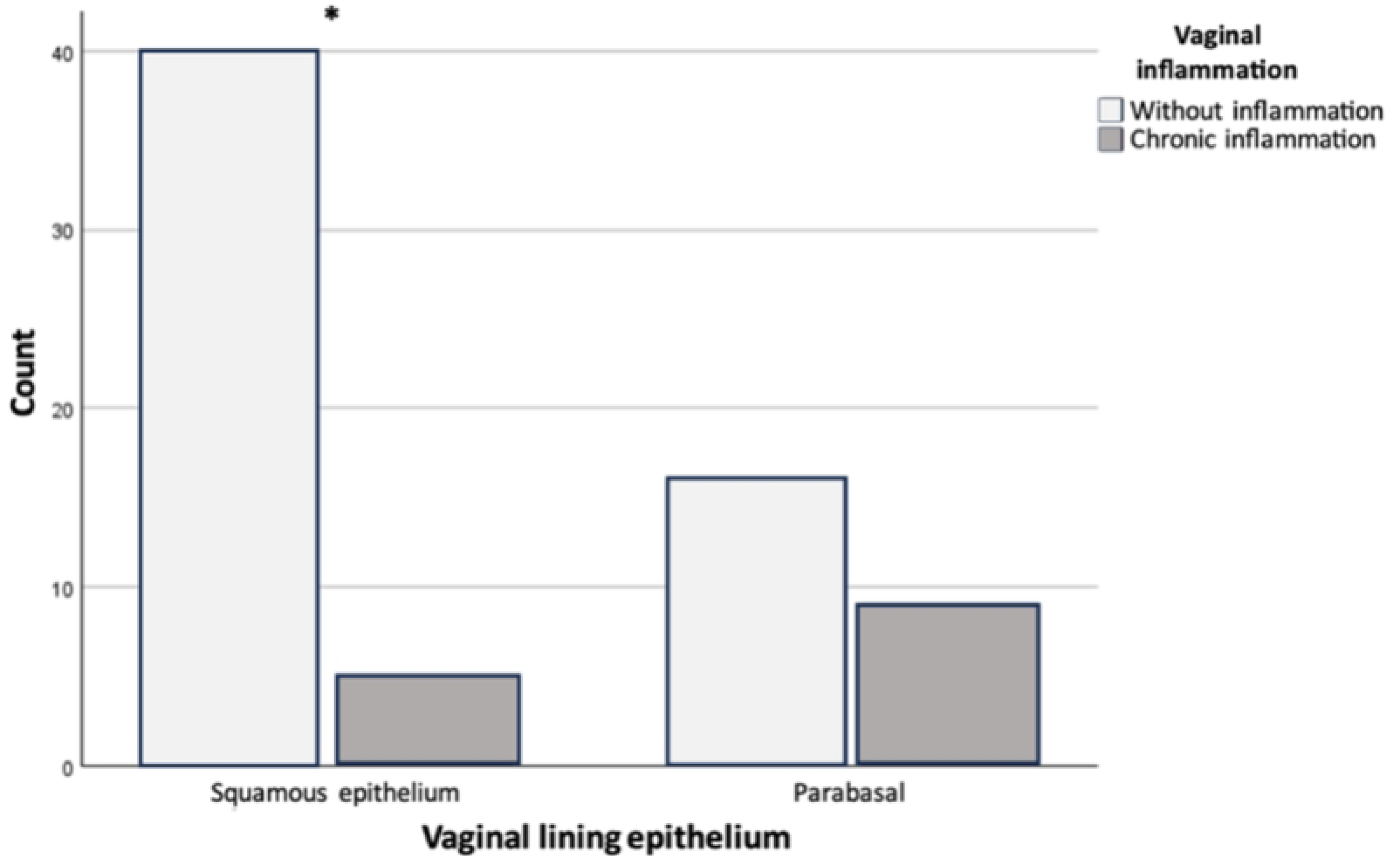
Relationship between vaginal lining epithelium and inflammation. *Asterisks (∗) indicate statistically significant differences (p < 0.05).* The chi-square test revealed a statistically significant association between epithelial type and inflammation (*p* = 0.013), indicating that squamous epithelium is more frequently associated with the absence of inflammation. **Note:** This analysis does not compare treatment groups, but rather categorical histological variables.

**Table 7:**
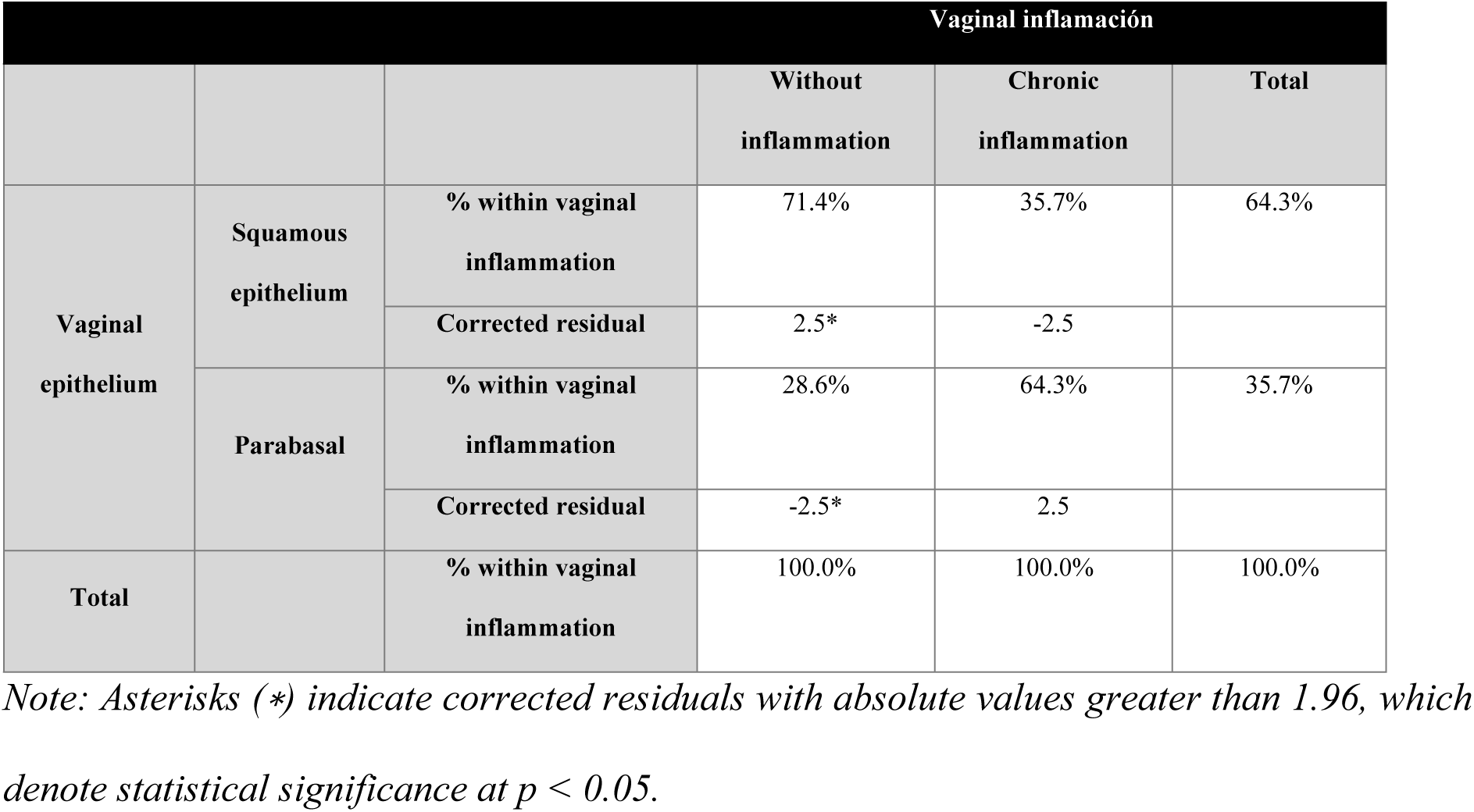
Vaginal lining epithelium and inflammation contingency table.

**Table 8:** Vaginal lining epithelium and vaginal inflammation Chi-square test.

|  | Value | gl | Asymptotic significance<br>(bilateral) | Exact significance<br>(bilateral) | Exact<br>significance<br>(unilateral) |
| --- | --- | --- | --- | --- | --- |
| Pearson's chi-square test | 6.222<sup>a</sup> | 1 | 0.013* |  |  |
| Continuity correction | 4.764 | 1 | 0.029 |  |  |
| Likelihood ratio | 5.991 | 1 | 0.014 |  |  |
| Fisher's exact test |  |  |  | 0.026 | 0.016 |

<sup>a</sup> Pearson Chi-Square test was used; $p < 0.05$ values are marked with an asterisk (\*) to indicate statistical significance.

**Table 9:**
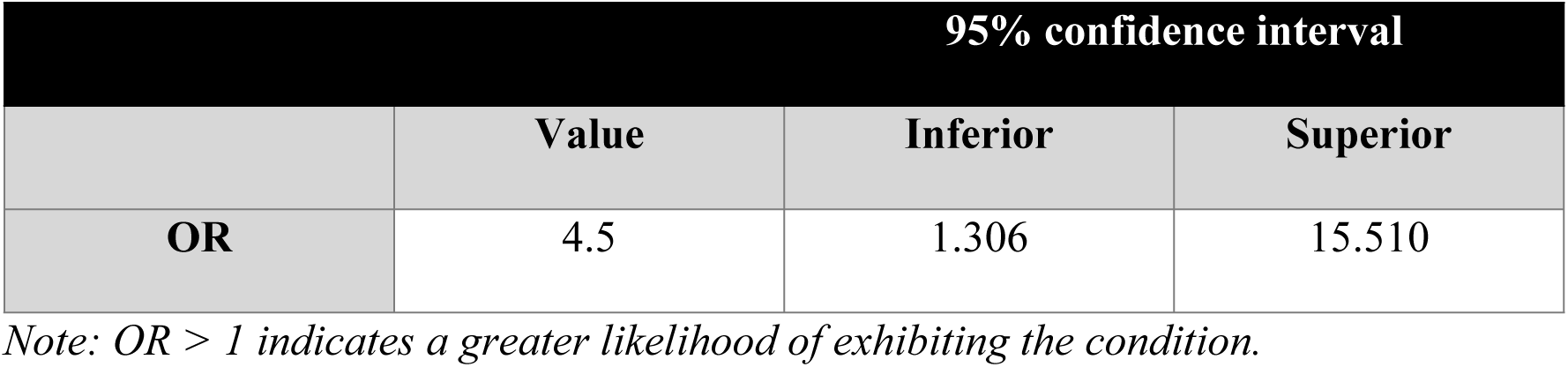
Odds ratio (OR) for Vaginal lining epithelium and vaginal inflammation.

#### Relationship Between Number of Epithelial Layers and Vaginal Inflammation

A significant association was found between the number of vaginal epithelial layers and the absence of inflammation (p < 0.001; Table 10). The contingency table revealed that when the epithelium contains seven or fewer layers, chronic inflammation is more frequent, indicating an inverse relationship. The most notable corrected residual was 4.6 for the 7–8 layer range, highlighting a strong association with chronic inflammation (Table 11). Fig 8 shows that inflammation was absent in all cases with nine or more epithelial layers. This association was further confirmed as statistically significant by Cramér’s V test (p = 0.001; Table 12).

**Fig 8.**
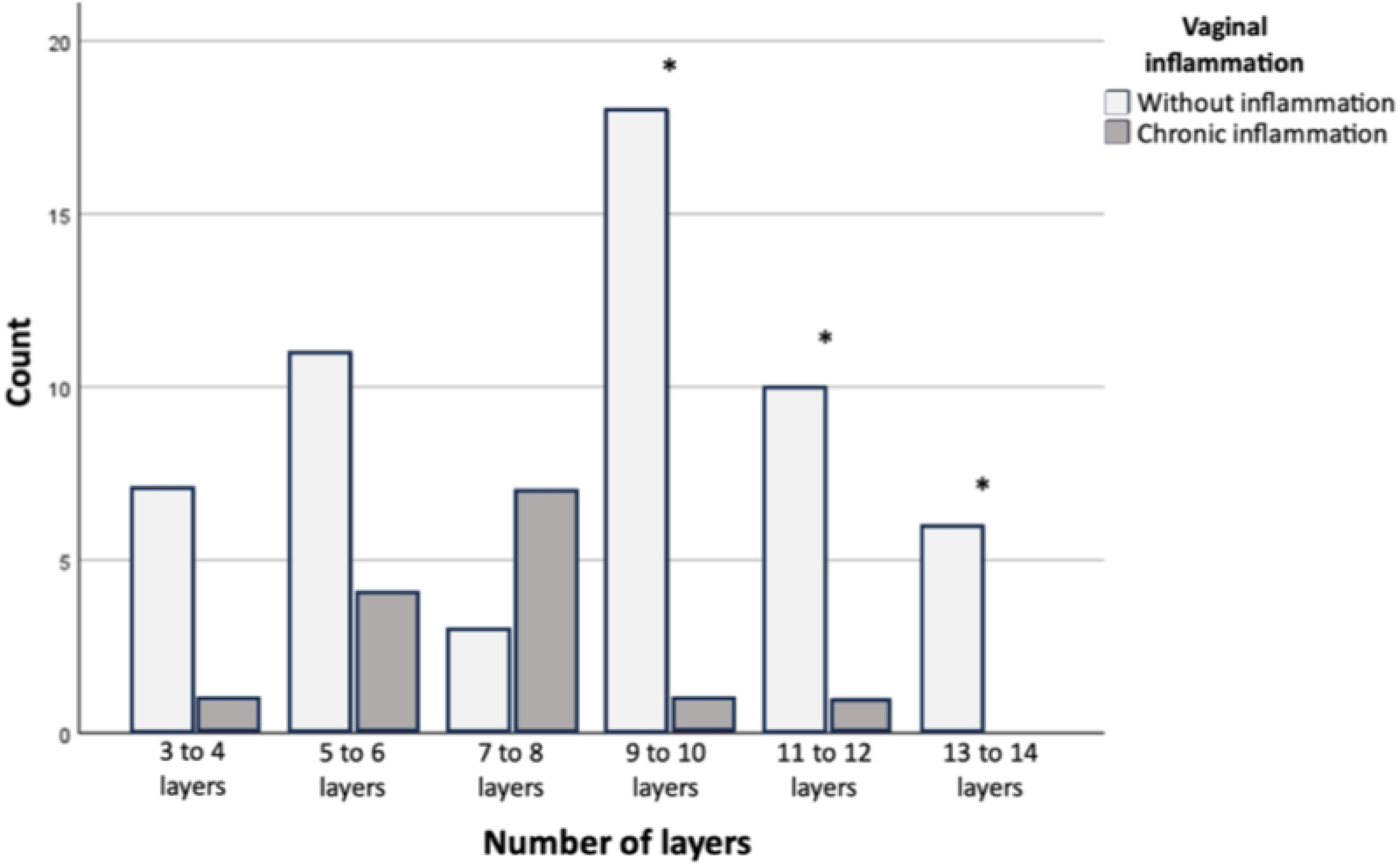
Relationship between number of epithelial layers and vaginal inflammation. Animals with nine or more epithelial layers showed no signs of chronic inflammation, while those with seven or fewer layers showed a higher frequency of inflammatory response. A statistically significant inverse relationship was observed between the number of layers and vaginal inflammation (*p* < 0.001, Pearson’s chi-square test). **(*)** indicates statistical significance. The figure compares categorical ranges, not experimental treatments.

**Table 10:** Contingency table of number of layers and Vaginal inflammation.

|  |  |  | Vaginal inflammation |  |  |
| --- | --- | --- | --- | --- | --- |
|  |  |  | Without inflammation | Chronic inflammation | Total |
| Number of layers | 3 to 4 layers | % within vaginal inflammation | 12.7% | 7.1% | 11.6% |
|  |  | Corrected residual | 0.6 | -0.6 |  |
|  | 5 to 6 layers | % within vaginal inflammation | 20.0% | 28.6% | 21.7% |
|  |  | Corrected residual | -0.7 | 0.7 |  |
|  | 7 to 8 layers | % within vaginal inflammation | 5.5% | 50.0% | 14.5% |
|  |  | Corrected residual | -4.2* | 4.2 |  |
|  | 9 to 10 layers | % within vaginal inflammation | 32.7% | 7.1% | 27.5% |
|  |  | Corrected residual | 1.9 | -1.9 |  |
|  | 11 to 12 layers | % within vaginal inflammation | 18.2% | 7.1% | 15.9% |
|  |  | Corrected residual | 1.0 | -1.0 |  |
|  | 13 to 14 layers | % within vaginal inflammation | 10.9% | 0.0% | 8.7% |
|  |  | Corrected residual | 1.3 | -1.3 |  |
| Total |  | % within vaginal inflammation | 100.0% | 100.0% | 100.0% |

*Note: Asterisks (\*) indicate corrected residuals with absolute values greater than 1.96, which* *denote statistical significance at $p < 0.05$ .*

**Table 11:** Chi-square of number of layers and Vaginal inflammation.

|  | Value | gl | Asymptotic significance<br>(bilateral) |
| --- | --- | --- | --- |
| Pearson's chi-square test | 20.989 <sup>a</sup> | 5 | < 0.001* |
| Likelihood ratio | 19.426 | 5 | 0.002 |
| Linear by linear association | 3.009 | 1 | 0.083 |
| Number of valid cases | 69 |  |  |

*Note: Chi-Square test was used; $p < 0.05$ value is marked with an asterisk (\*) to indicate* *statistical significance.*
<sup>a</sup> *Seven cells (58.3%) have expected count less than 5. The minimum expected count is 1.22.*

#### Relationship Between Number of Layers and Vaginal Epithelial Lining

The contingency table (Table 13) shows that when the number of epithelial layers is low (7–8 layers), squamous epithelium is largely absent, with parabasal flat epithelium being the predominant type. Among cases with 9–10 layers, 40.9% exhibited squamous epithelium, while 25% of those with 11–12 layers also showed this epithelial type. This association was statistically significant (p = 0.001; Table 14) and demonstrated a strong correlation, as indicated by a high Cramér’s V value of 0.786 (Table 15). Fig 9 illustrates that parabasal flat epithelium predominates in tissues with fewer than eight layers, whereas squamous epithelium becomes increasingly predominant as the number of layers increases.

**Fig 9.**
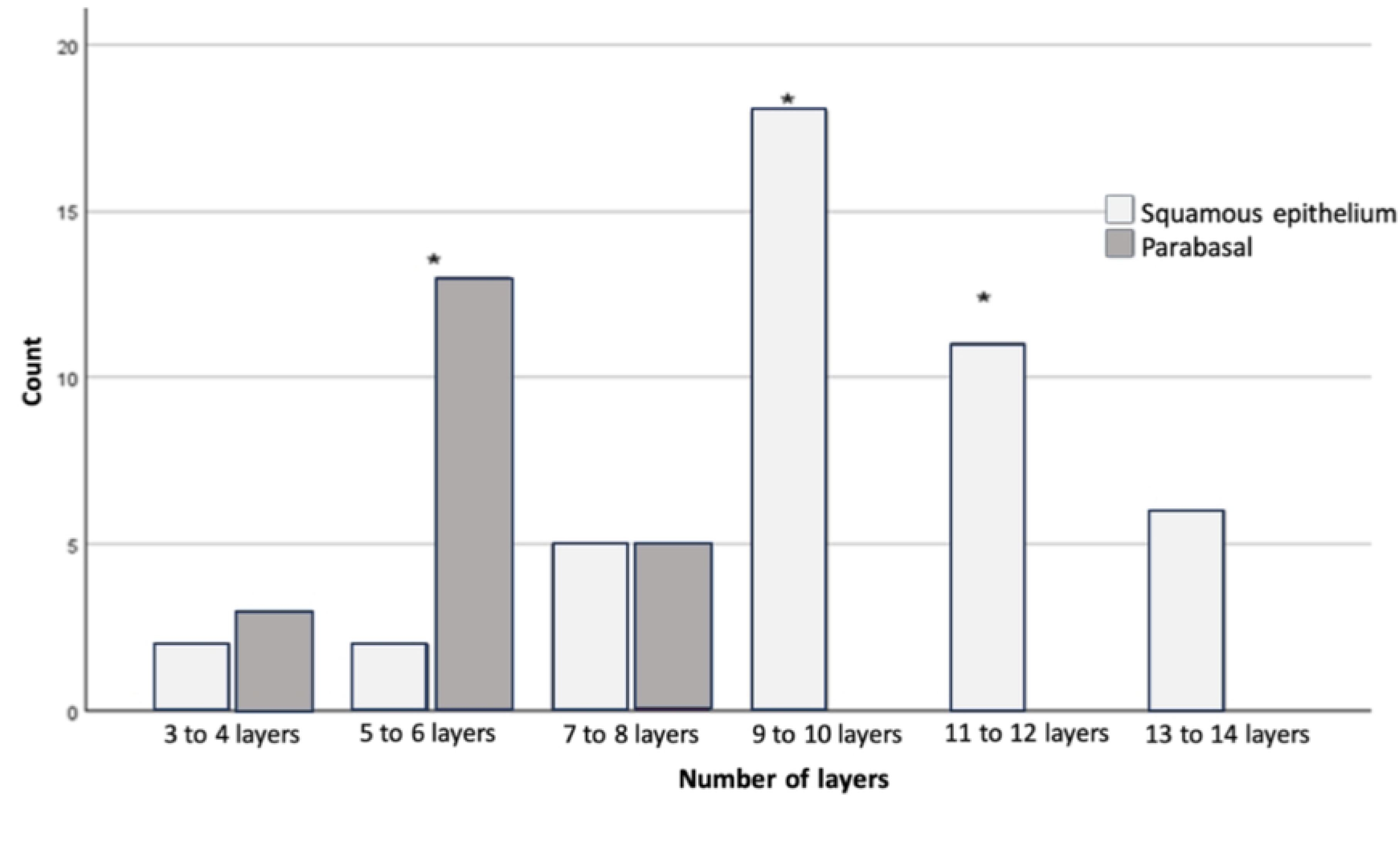
Relationship between number of layers and vaginal lining epithelium. A significant association was found between lower numbers of epithelial layers (≤6) and the presence of parabasal epithelium, while nine or more layers were associated with the presence of squamous epithelium (*p* < 0.001). Corrected residual analysis showed that parabasal epithelium was significantly more frequent in tissues with 5–6 layers, whereas squamous epithelium predominated in tissues with 9–14 layers. **(*)** indicates statistically significant associations between epithelial type and layer count.

**Table 13:**
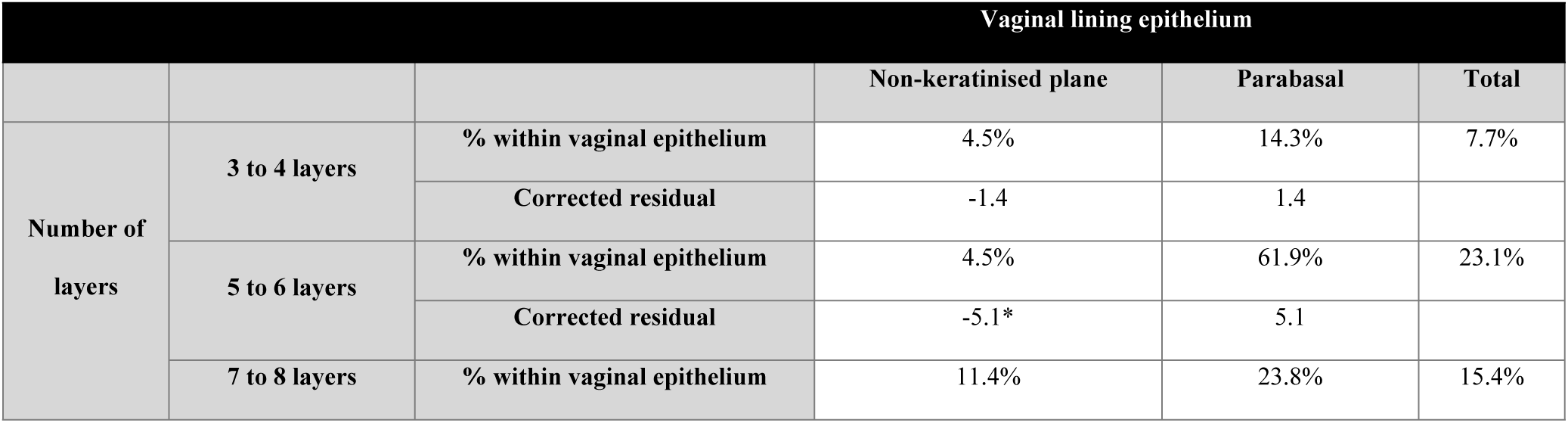
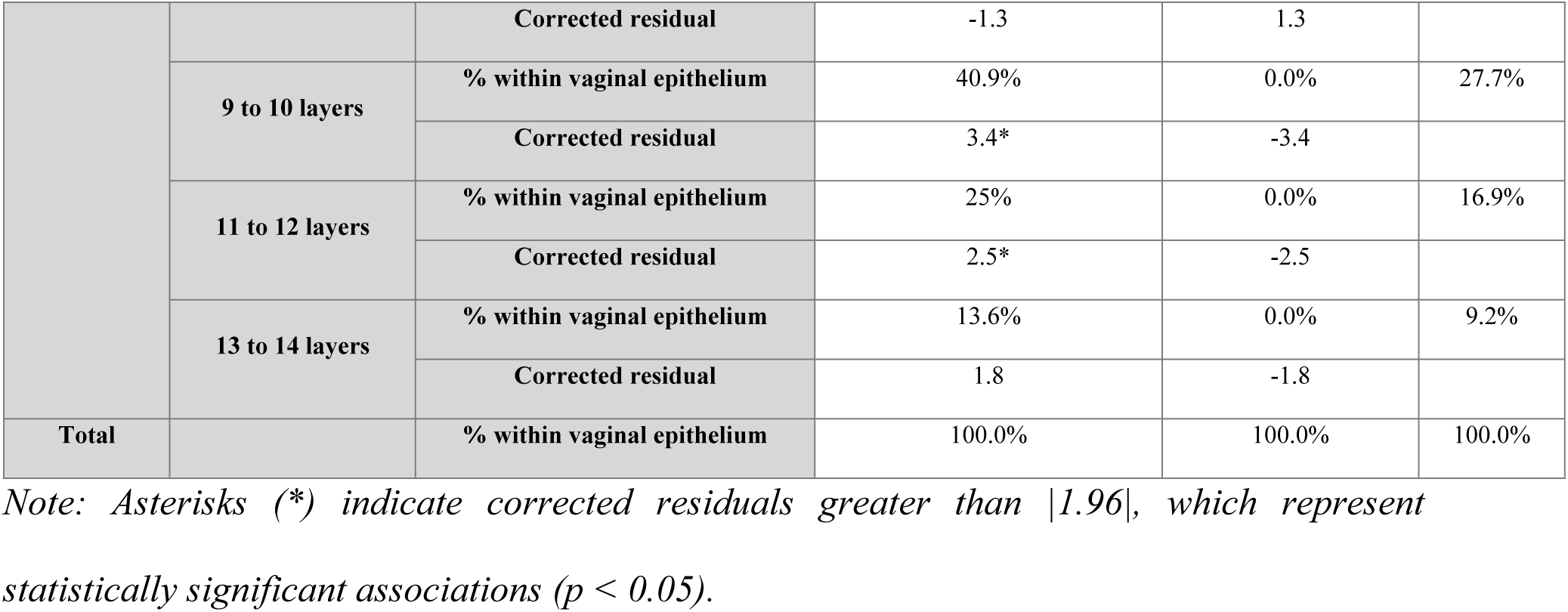
Contingency table of number of layers and Vaginal Lining Epithelium.

**Table 14:**
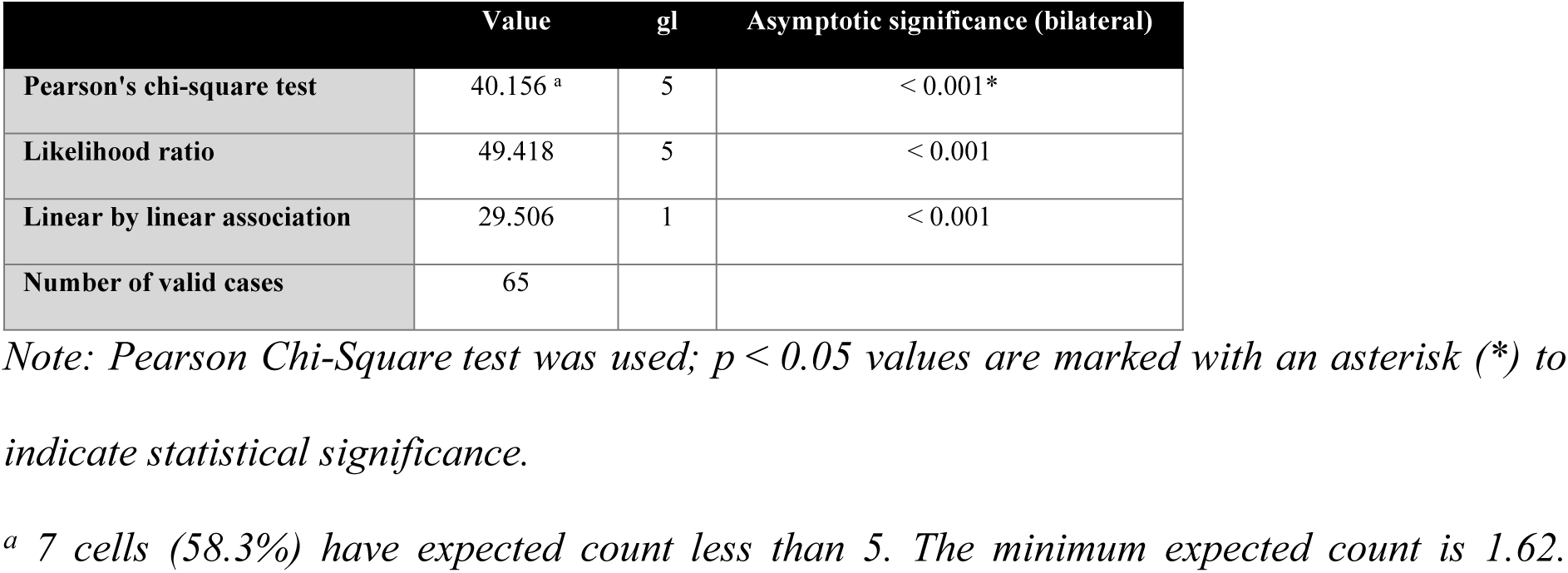
Chi-square of number of layers and Vaginal Lining Epithelium.

|  | Value | gl | Asymptotic significance (bilateral) |
| --- | --- | --- | --- |
| Pearson's chi-square test | 40.156 <sup>a</sup> | 5 | < 0.001* |
| Likelihood ratio | 49.418 | 5 | < 0.001 |
| Linear by linear association | 29.506 | 1 | < 0.001 |
| Number of valid cases | 65 |  |  |

Note: Pearson Chi-Square test was used; $p < 0.05$ values are marked with an asterisk (\*) to indicate statistical significance.
<sup>a</sup> 7 cells (58.3%) have expected count less than 5. The minimum expected count is 1.62.

**Table 15:** Cramer’s V-test of the relationship between number of layers and Vaginal lining epithelium.

|  |  | Value | Approximate significance |
| --- | --- | --- | --- |
| Nominal by nominal | Phi | 0.786 | < 0.001 |
|  | Cramér's V | 0.786 | < 0.001* |
| Number of valid cases |  | 65 |  |

Note: Asterisks (\*) indicates $p < 0.05$ indicates statistical significance according to Cramér's $V$ test.

#### Relationship of Vaginal Epithelial Lining with Number of Layers and Vaginal Inflammation

According to the contingency table (Table 16), specimens with squamous epithelium and more than nine epithelial layers exhibited no signs of vaginal inflammation. Specifically, 45% of cases with 9–10 layers showed both squamous epithelium and absence of inflammation. In contrast, no specimens with parabasal flat epithelium had more than nine layers or were free of inflammation (Table 16, Fig 10). The chi-square test revealed a significant association (p < 0.001), indicating a relationship between the absence of inflammation, the presence of squamous epithelium, and having more than nine epithelial layers (Table 17). This relationship was further supported by a strong and statistically significant Cramér’s V value of 0.811 (p < 0.001; Table 18).

**Fig 10.**
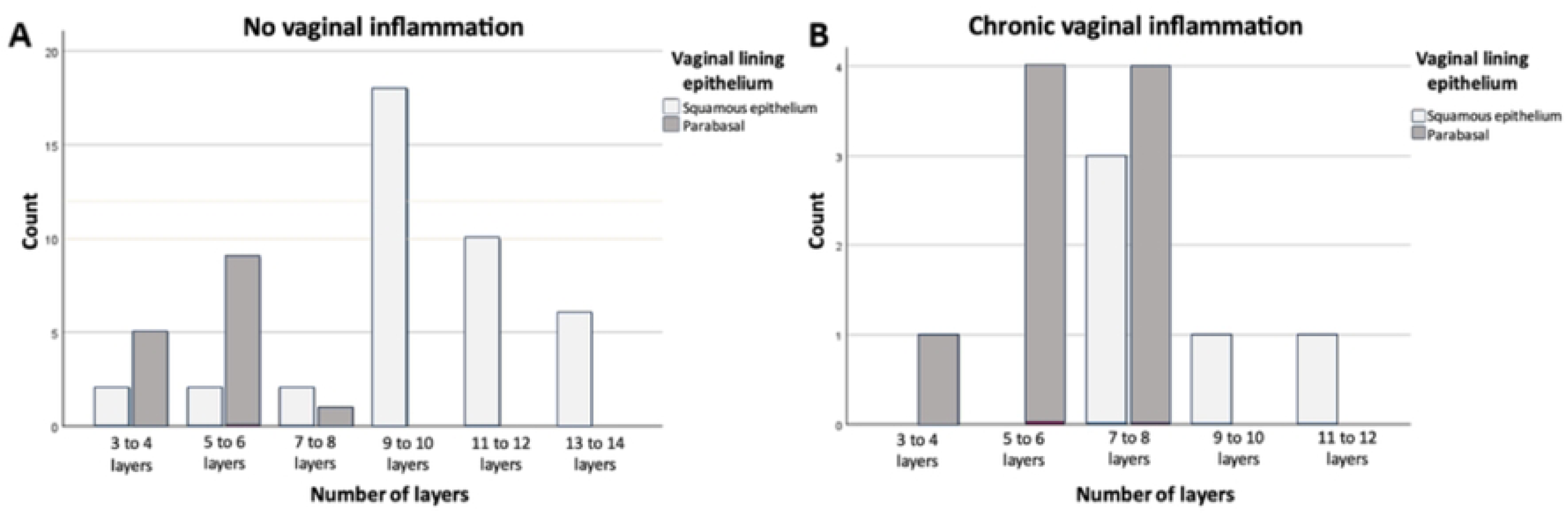
Combined relationship between vaginal epithelial type, number of epithelial layers, and inflammation status. The contingency analysis shows that specimens with squamous epithelium and more than nine epithelial layers exhibited no signs of vaginal inflammation. Conversely, tissues with parabasal epithelium and fewer than seven layers were more frequently associated with chronic inflammation. This three-way association was statistically significant according to the chi-square test (*p* < 0.001), and supported by a strong effect size (Cramér’s V = 0.811). **Note:** No individual comparisons were marked with asterisks in the figure.

**Table 16:**
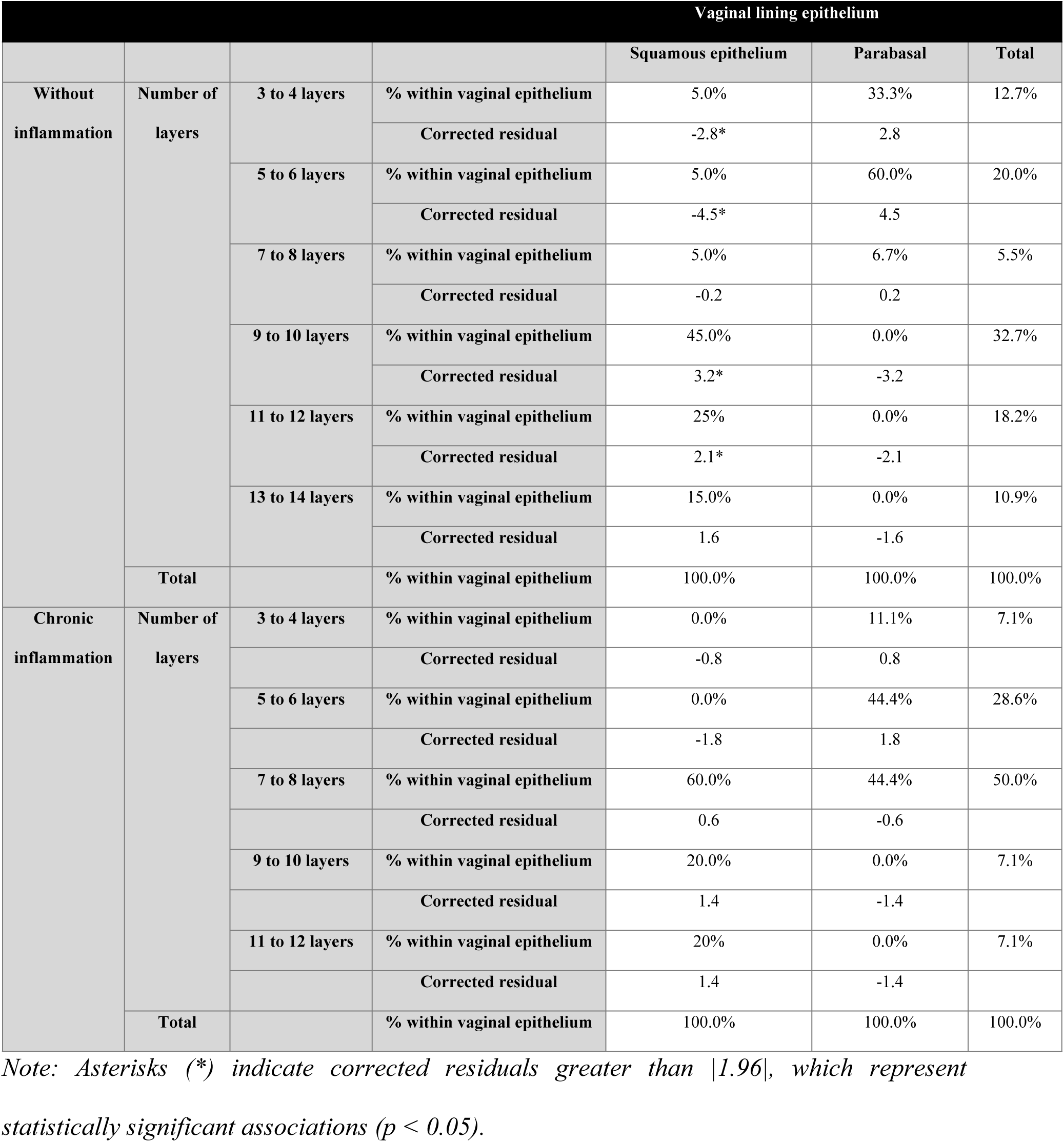
Contingency table of Vaginal Lining Epithelium, number of layers and vaginal inflammation.

**Table 17:** Chi-square of Vaginal lining epithelium, number of layers and vaginal inflammation.

|  |  | Value | gl | Asymptotic significance<br>(bilateral) |
| --- | --- | --- | --- | --- |
| Without inflammation | Pearson's chi-square test | 36.187 <sup>a</sup> | 5 | < 0.001* |
|  | Likelihood ratio | 41.829 | 5 | < 0.001 |
|  | Linear by linear association | 27.942 | 1 | < 0.001 |
|  | Number of valid cases | 55 |  |  |
| Chronic inflammation | Pearson's chi-square test | 6.533 <sup>b</sup> | 4 | 0.163 |
|  | Likelihood ratio | 8.688 | 4 | 0.069 |
|  | Linear by linear association | 5.425 | 1 | 0.020 |
|  | Number of valid cases | 14 |  |  |
| Total | Pearson's chi-square test | 43.726 <sup>c</sup> | 5 | < 0.001 |
|  | Likelihood ratio | 54.520 | 5 | < 0.001 |
|  | Linear by linear association | 34.349 | 1 | < 0.001 |
|  | Number of valid cases | 69 |  |  |

Note: Asterisk (\*) indicates statistical significance ( $p < 0.05$ ) based on the Pearson chi-square test. When more than 20% of expected frequencies are below 5, results should be interpreted with caution, as chi-square assumptions may be violated.
<sup>a</sup> 8 cells (66.7%) have expected counts less than 5. The minimum expected count is 0.82.
<sup>b</sup> 10 cells (100.0%) have expected counts less than 5. The minimum expected count is 0.36.
<sup>c</sup> 5 cells (41.7%) have expected counts less than 5. The minimum expected count is 2.09.

**Table 18:**
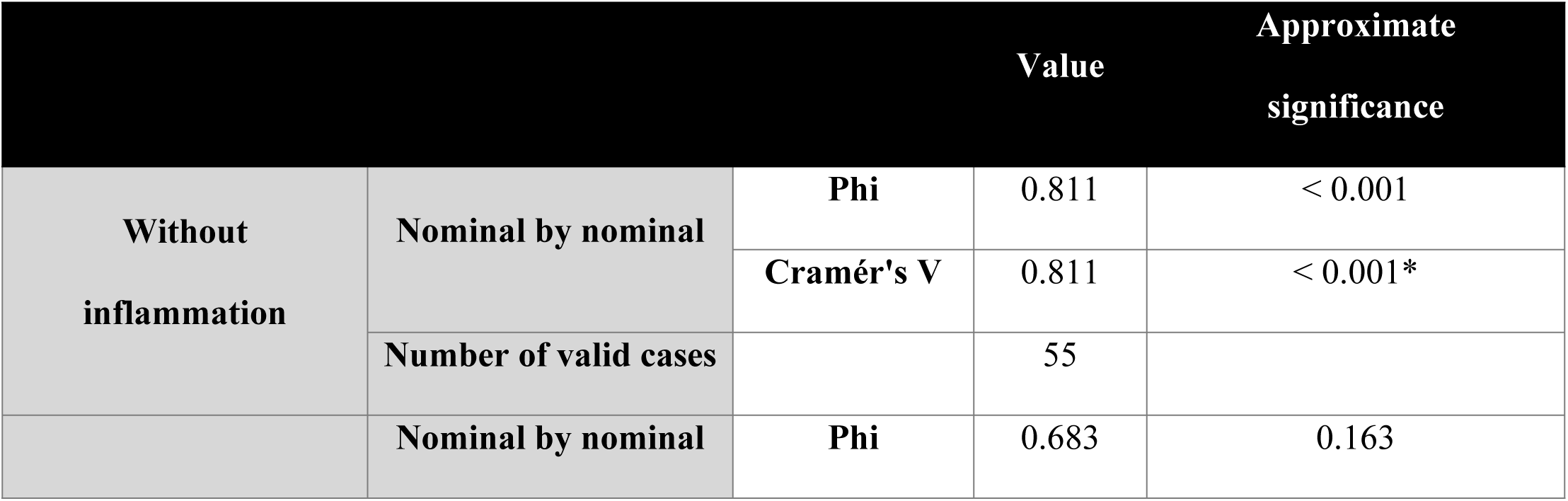
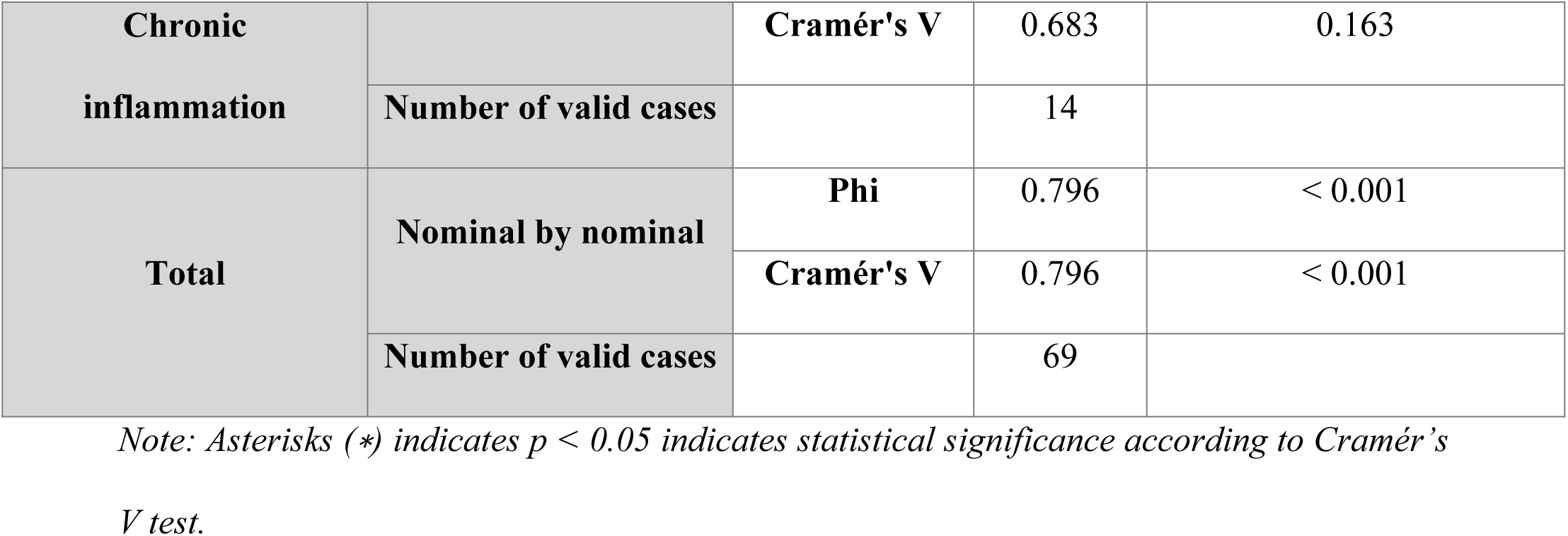
Cramer’s V test of the relationship between vaginal lining epithelium, number of layers and vaginal inflammation.

### Weight Analysis

#### Vaginal Weight

Compared to the control group (C0), the PL20% and PL20%-UG groups showed significantly lower vaginal weights (p = 0.038 and p = 0.020, respectively), while no significant differences were observed in the remaining groups (Fig 11). The Kolmogorov–Smirnov test indicated a normal distribution of vaginal weight data (p = 0.068). However, Levene’s test revealed heterogeneity of variances among groups (p = 0.018), prompting the use of Welch’s ANOVA, which confirmed a significant difference in mean vaginal weight between groups (p = 0.013; Table 19). Subsequent chi-square comparisons (Table 20) revealed that the PL20% and PL20%-UG groups had statistically significant differences compared to C0 (*p* < 0.05).

**Fig 11.**
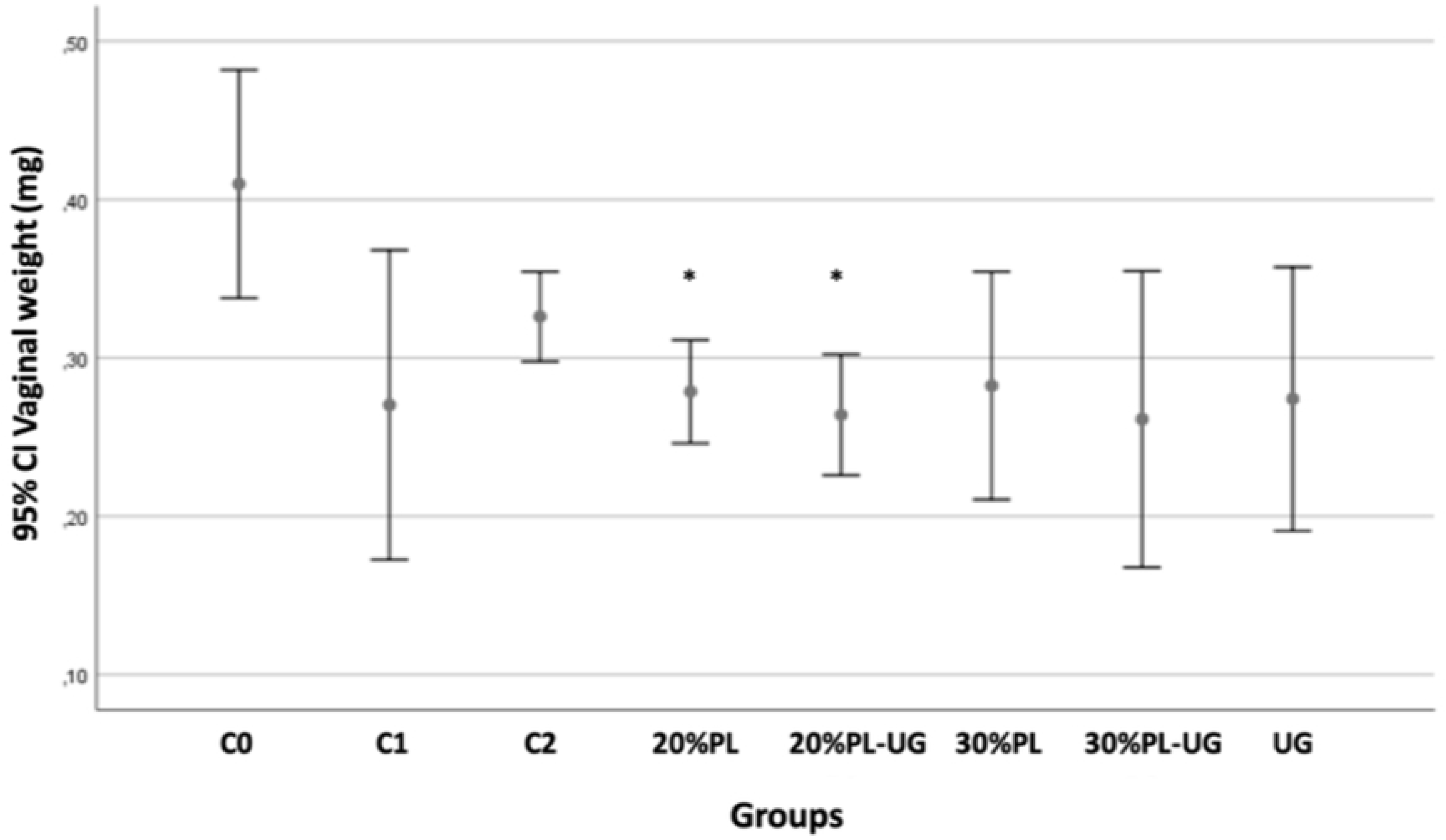
Vaginal weight diagram. C0: Control without surgery; C1: Control without treatment; C2: Control with estrogen. *Asterisks (∗) indicate statistically significant differences compared to C0 (p < 0.005)*.

**Table 19:** Welch’s Vaginal Weight Test.

| Robust tests of equality of means |  |  |  |  |
| --- | --- | --- | --- | --- |
|  | Statistic <sup>a</sup> | gl1 | gl2 | Sig. |
| **Welch** | 3.216 | 7 | 26.234 | 0.013 |

*Note: Welch's ANOVA was applied due to variance heterogeneity detected by Levene's test ( $p$* *= 0.018). The test revealed a significant difference in mean vaginal weight among groups ( $p$ =* *0.013).*
<sup>a</sup> *Asymptotically F-distributed.*

**Table 20:** Comparison of C0 means with the other Vaginal Weight groups.

|  | C1 |  | C2 |  | 20% PL |  | 20% PL-UG |  | 30% PL |  | 30% PL-UG |  | UG |  |
| --- | --- | --- | --- | --- | --- | --- | --- | --- | --- | --- | --- | --- | --- | --- |
|  | DME ± SE | Sig | DME<br>± SE | Sig | DME ±<br>SE | Sig | DME ±<br>SE | Sig | DME ± SE | Sig | DME ± SE | Sig | DME ± SE | Sig |
| **C0** | 0.14 ± 0.053 | 0.217 | 0.08 ±<br>0.34 | 0.299 | 0.13 ±<br>0.035 | 0.038* | 0.15 ± 0.03 | 0.020* | 0.12 ± 0.044 | 0.144 | 0.14 ± 0.05 | 0.143 | 0.135 ±<br>0.047 | 0.151 |

*Note: C0 = no inflammation; C2 = chronic inflammation; PL = Platelet lysate; UG = Uncaria*
*tomentosa; SMD = standardized mean difference; SE = standard error.*
*Asterisks (\*) indicate statistically significant differences compared to C0 ( $p < 0.05$ ), based on* *the chi-square test.*

#### Uterine Weight / Body Weight Ratio

As shown in Fig 12, group C0 exhibited a median uterine weight/body weight ratio of 0.0022 × 10⁻³, with greater dispersion compared to other groups. In contrast, the PL20% and PL20%-UG groups had lower median values of 0.0004 × 10⁻³ and 0.00045 × 10⁻³, respectively. A statistically significant reduction in this ratio was observed in groups C1 (p = 0.030), PL20%-UG (p = 0.005), PL30% (p = 0.026), and UG (p = 0.010) compared to C0 (Table 21).

**Fig 12.**
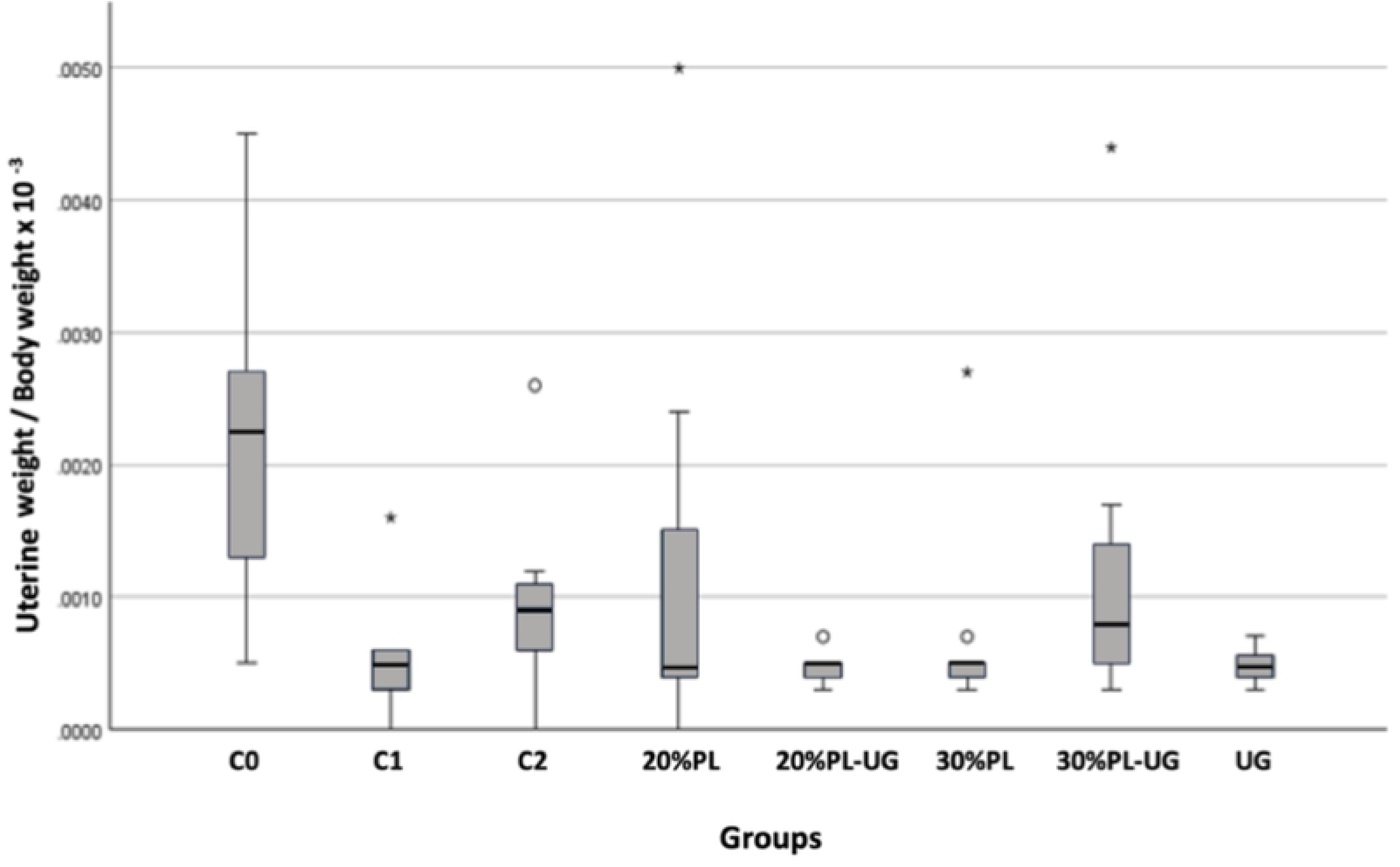
Comparison of uterine weight to body weight ratio. C0: Control without surgery; C1: Control without treatment; C2: Control with estrogen. *Asterisks (∗) indicate statistically significant differences compared to C0 (p < 0.005)*.

**Table 21:** Comparison of averages in pairs.

|  | C1 | C2 | 20% PL | 20% PL-UG | 30% PL | 30% PL-UG | UG |
| --- | --- | --- | --- | --- | --- | --- | --- |
| C0 | 0.030* | 1.000 | 0.219 | 0.005* | 0.026* | 1.000 | 0.010* |

*Note: C0 = no inflammation; C2 = chronic; PL = Platelet lysate; UG = Uncaria tomentosa.*
*Asterisks (\*) indicate statistically significant differences according to the Kruskal–Wallis test*
*( $p < 0.05$ ).*

#### Vaginal Weight / Body Weight Ratio

Compared to group C0, a significant reduction in the vaginal weight/body weight ratio was observed in groups C1 (p = 0.004), PL20% (p = 0.021), PL30% (p = 0.045), and PL30%-UG (p = 0.008) (Fig 13) (Table 22).

**Fig 13.**
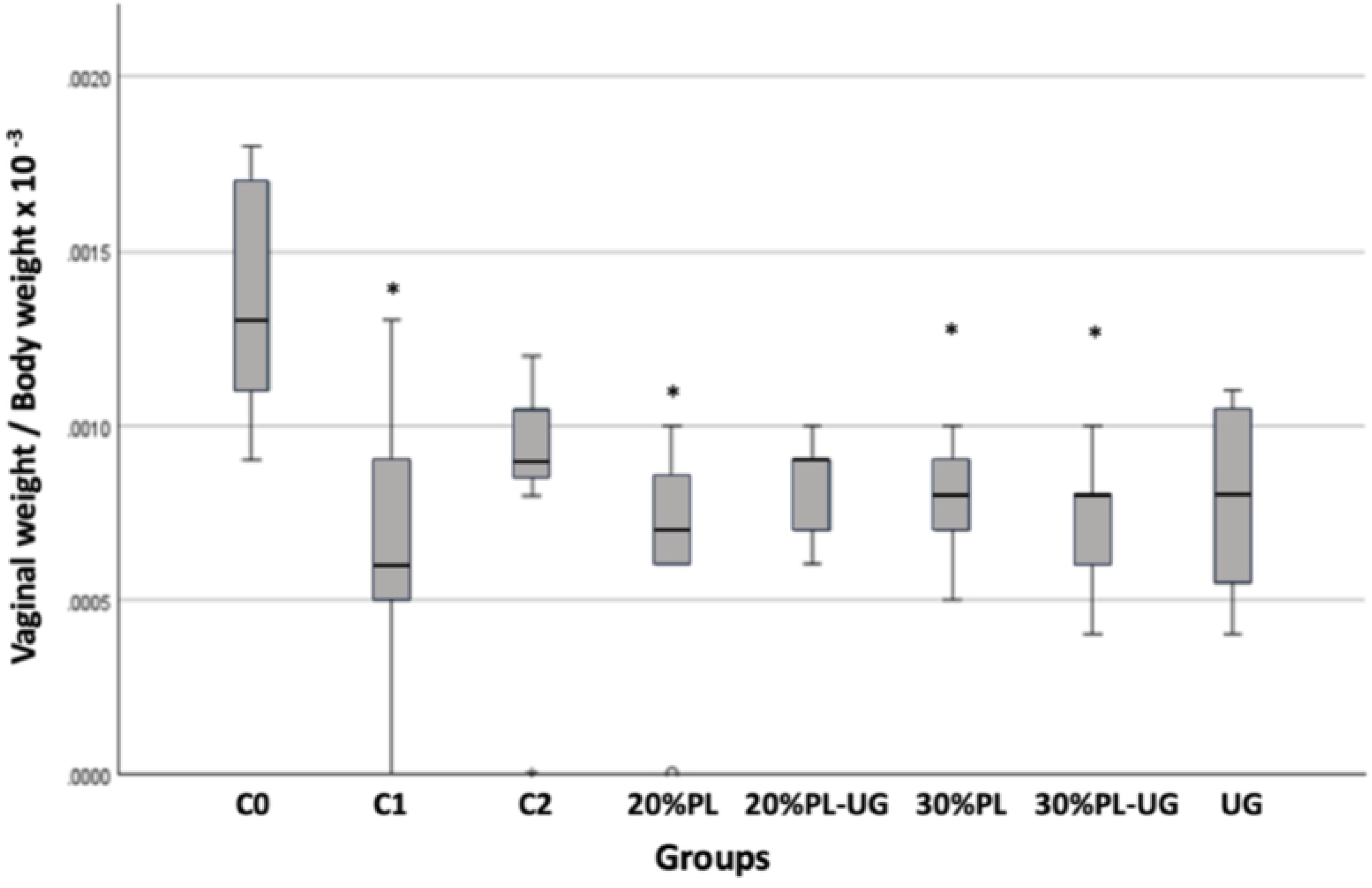
Diagram of vaginal weight relative to body weight. C0: Control without surgery; C1: Control without treatment; C2: Control with estrogen. *Asterisks (∗) indicate statistically significant differences compared to C0 (p < 0.005)*.

**Table 22:**
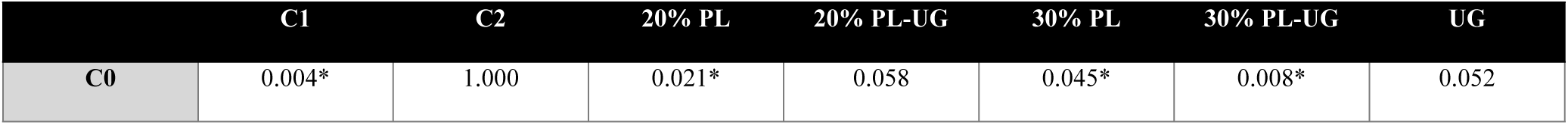
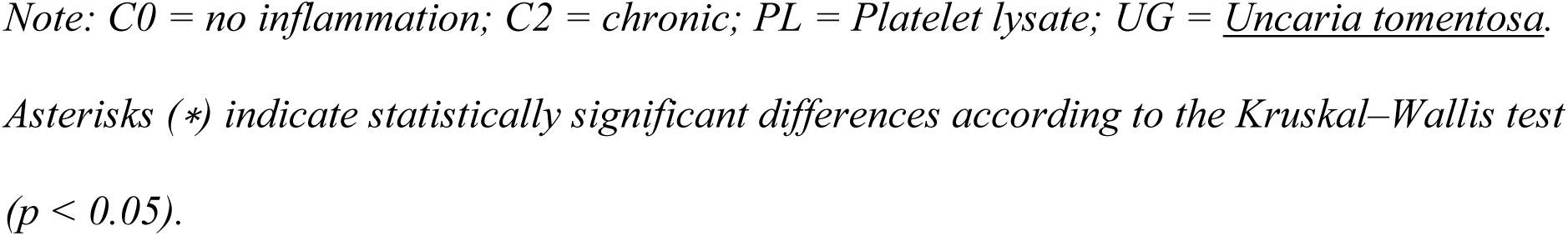
Comparison of averages in pairs.

|  | C1 | C2 | 20% PL | 20% PL-UG | 30% PL | 30% PL-UG | UG |
| --- | --- | --- | --- | --- | --- | --- | --- |
| C0 | 0.004* | 1.000 | 0.021* | 0.058 | 0.045* | 0.008* | 0.052 |

*Note: C0 = no inflammation; C2 = chronic; PL = Platelet lysate; UG = Uncaria tomentosa.* *Asterisks (\*) indicate statistically significant differences according to the Kruskal–Wallis test* *( $p < 0.05$ ).*

## Discussion

The estrous cycle of the rats was confirmed previous ovariectomy by vaginal cytology, identifying characteristic cell types and their proportions. Following ovariectomy, the low number of squamous cells indicated the absence of estrus (25), evidencing a lack of estrogen production. This was further confirmed by the ELISA kit ELK8714, which showed a significant decrease in estradiol levels in all operated groups except the control group C0. These, cytology and estradiol deficiency confirmed the establishment of a postmenopausal rat model (16)

This study compared different treatments with natural products aimed at repairing and reducing inflammation caused by vaginal atrophy in the absence of hormones. Macroscopically, no significant differences were observed in vulvar coloration, which remained red in all groups, nor in uterine thickness, which was thin in the treated groups—findings consistent with those reported by (26)

Although the PL30% group exhibited greater vaginal inflammation than other groups, this difference was not statistically significant. High concentrations of PL may explain to induce an inflammatory environment, as the growth factors present promote granulation tissue formation and proinflammatory cytokine release (26). The group treated with cat’s claw extract (UG) showed no significant inflammation compared to controls, possibly due to its high phenolic content that inhibits proinflammatory mediators; however, this effect was diminished when UG was combined with PL30%.

Regarding the vaginal epithelial lining, the UG group showed a significant decrease in squamous epithelium with predominance of parabasal cells. Conversely, treatments with PL20%, PL20%-UG, and PL30% favored the presence of squamous epithelium, similar to tissue repair observed in the estrogen-treated group and in studies using dehydroepiandrosterone, which has estrogenic and androgenic activity and reverses the loss of squamous epithelium (27). PL20% promoted an epithelium of 3 to 6 layers, while PL30% and PL20%-UG achieved 9 to 10 layers, comparable to controls, without the need for steroid hormones, which reverse epithelial thinning caused by vaginal atrophy (28). In contrast, UG presented an epithelium of 5 to 6 layers, and PL30%-UG showed a broad range of 3 to 10 layers.

A significant relationship was observed between the predominance of squamous epithelium and the absence of inflammation, with a 4.5 times higher probability of no inflammation in the presence of squamous epithelium compared to parabasal epithelium. Furthermore, the number of epithelial layers was inversely related to chronic inflammation and directly related to the presence of squamous epithelium. Groups PL20%, PL20%-UG, and PL30% showed greater squamous epithelial repair, higher number of layers, and absence of inflammation, similar to controls C0 and C2. Conversely, PL30%-UG presented significant inflammation and fewer layers, and UG failed to restore squamous epithelium, exhibiting only parabasal epithelium with fewer layers. This suggests that chronic inflammation in PL30%-UG impairs epithelial repair by affecting collagen remodeling, promoting fibrosis, and altering tissue regeneration (29,30).

Regarding weights, the C1 group (ovariectomy without treatment) showed significantly lower uterine weight than C0, reflecting uterine atrophy due to estrogen deficiency, with decreased endometrial and myometrial thickness (28), similar to groups treated with PL30%, UG, and PL20%-UG. In contrast, PL20% and PL30%-UG did not differ significantly from C0, indicating preservation of uterine thickness. Treatment with 0.005% estradiol (C2) showed lower uterotrophic activity, without increased myometrial thickness (31).

## Conclusion

Treatments with 20% PL, with or without cat’s claw extract (UG), showed favorable results similar to the estradiol-treated group, without presenting side effects. Treatment with 30% PL induced inflammation, which was significant when combined with UG, affecting its anti-inflammatory effect. The PL30%-UG group did not show significant tissue repair due to chronic inflammation inhibiting epithelial regeneration. There is a significant relationship between squamous epithelium repair, absence of inflammation, and increased number of vaginal epithelial layers. Groups treated with 20% PL, with or without UG, and 30% PL showed results comparable to hormonal treatments, without uterotrophic effects, as none significantly increased uterine weight compared to the unoperated control group.

Future studies are recommended to include the measurement of pro-inflammatory and anti-inflammatory cytokines, as well as hormone levels at both serum and local tissue levels. It is suggested that new research focuses on the evaluation of uterine atrophy to complement the vaginal findings.

The sample size used (n = 9 per group) was based on ethical and methodological considerations; however, it represents a limitation of the study. Future research should replicate these findings using larger sample sizes to improve statistical power and reproducibility.

Importantly, it would be valuable to explore the clinical implications and potential therapeutic use of PL and cat’s claw extract could offer a novel, non-hormonal therapeutic approach for treating postmenopausal vaginal atrophy. Therefore, these findings warrant further investigation through preclinical safety studies and eventual clinical trials in postmenopausal women to assess efficacy, tolerability, and long-term outcomes.

## Acknowledgements

We would like to thank the Universidad Peruana Cayetano Heredia for institutional support during the development of this project.

